# Diffraction minima resolve point scatterers at tiny fractions (1/80) of the wavelength

**DOI:** 10.1101/2024.01.24.576982

**Authors:** Thomas A. Hensel, Jan O. Wirth, Stefan W. Hell

## Abstract

Discerning two or more identical and constantly scattering point sources using freely propagating waves is thought to be limited by diffraction. Here we show both theoretically and experimentally that by employing a diffraction minimum rather than a maximum for resolution, a given number of point scatterers can be discerned at tiny fractions of the employed wavelength. Specifically, we identify an 8 nm distance between two constantly emitting (non-blinking, non-switchable) fluorescent molecules, corresponding to 1/80 of the wavelength. Moreover, we show that contrary to naïve expectations, the measurement precision improves with decreasing distance between the scatterers and with increased scatterer density, thus opening up the prospect of resolving clusters of (optical) point scatterers at tiny fractions of the wavelength.

---

A prominent question in physics is how small the distance *d* between two simultaneously scattering point sources can be in order to still be separable by focused (light) waves of wavelength *λ*. Separating two inelastic optical scatterers and measuring their distance is important in many fields, particularly in fluorescence microscopy, which is the most widely applied imaging modality in the life sciences. Fluorescent molecules, also known as fluorophores, can be treated as inelastic optical scatterers, because by dissipating a part of the absorbed photon energy and emitting a photon of longer wavelength, these molecules annihilate any phase information of the incoming light field. In any case, the textbook answer to the question of the minimally resolvable distance is the Rayleigh criterion^1,2^. It states that the distance between the diffraction maxima of the scatterers in an image should not be smaller than *d* = 0.61 *λ*/*(n sin α)*, with *n* and *sin α* denoting the refractive index of the immersion medium and the sine of the half-aperture angle of the lens, respectively. While introduced for widefield microscopy, Rayleigh’s limit also applies to popular scanning optical microscopy, where the object is scanned with a focused beam of wavelength *λ* and the image given by the number of photons registered per scanning position^3^.

Unlike many other microscopy methods, fluorescence microscopy does not map out the molecules of interest per se, such as the proteins in a cell, but the fluorophores that have been linked to those molecules as labels. Although disadvantageous at first glance, this aspect is critical in superresolution fluorescence microscopy or nanoscopy because to obtain subdiffraction resolution, these methods manipulate fluorophore states. Specifically, fluorophores that reside closer than the diffraction barrier are discerned by transiently transferring a fraction of them into a molecular state that emits photons (ON), whereas the remaining fraction is kept in a state that does not (OFF). Placing fluorophores in different states for a brief period of detection instantly provides separability, rendering separation by focusing to a tiny spot obsolete^4,5^. This ON/OFF state transition is a key physical element of all diffraction-unlimited fluorescence microscopy methods known to date, including those called STED, PALM/STORM, and the more recently introduced approaches called MINFLUX^6^ and DNA-PAINT^7^. If this element were to be removed, none of them could provide nanoscale resolution.

Unfortunately, the separation by ON/OFF molecular states comes with fundamental limitations that preclude extending superresolution beyond fluorescence, e.g. to other types of (inelastic) optical scattering such as Raman, let alone to other areas of physics resolving with diffracted waves. Even within the realm of fluorescence microscopy, the ON/OFF principle inevitably demands that neighboring fluorophores must be recorded sequentially. This drawback is intolerable if adjacent fluorophores need to be observed and resolved at the same time, a situation that is commonly encountered in biophysical experiments involving molecular tracking. Hence, motivated by both fundamental and practical reasons, as well as the recent success of MINFLUX^6^, we reinvestigated the ability of optical microscopy to resolve individual scatterers that remain identical throughout the observation process.

Here we show that scanning with a focused light field having a (central) diffraction minimum – rather than a maximum – allows to measure the distances between a known number of optically identical fluorophores down to single-digit nanometers. By measuring the positions of two fluorophores situated only 8 nm (*λ*/80)apart, we show that Rayleigh’s criterion grossly overestimates the minimal distance at which two inelastic scatterers can be resolved in practice. The underlying physical reason is that probing the scatterers with a diffraction intensity minimum modulates the scattered signal without modulating their state, whilst precluding the modulation from being drowned by noise. As a result, the continuous tracking of (at least) two fluorophores at nanometer distances becomes viable.

We also show that for a given signal to noise ratio (SNR) and background, the precision in separability in fact increases with decreasing distance between the scatterers. Defying naïve expectations, this arguably surprising characteristic also allows us to extend the method to a larger number of scatterers, which holds great potential for the optical investigation of dynamic (re)arrangements of proteins and other molecules with conventional optics at the nanoscale.

## Results and discussion

The diffraction maxima representing two fluorophores located at distance *d*<*λ*/2 overlap considerably in the image of an optical microscope. This holds both for those formed by diffracted fluorescence photons on a camera, as well as for the maxima gained by scanning a focused excitation beam across the focal plane and registering the fluorescence photons with a confocal point detector. In both cases, the identification of each individual fluorophore is heavily contingent upon the noise in the image, which is usually Poissonian (Figure 1, a,b). If the signal-to-noise ratio (SNR) was infinite and the point-spread-function (PSF) of the imaging system perfectly known, the fluorophores could always be resolved by deconvolution with the PSF. In practice, however, the knowledge of the PSF is compromised by aberrations and the SNR is by far too low for resolving the fluorophores reliably^8,9^.

**Figure 1:**
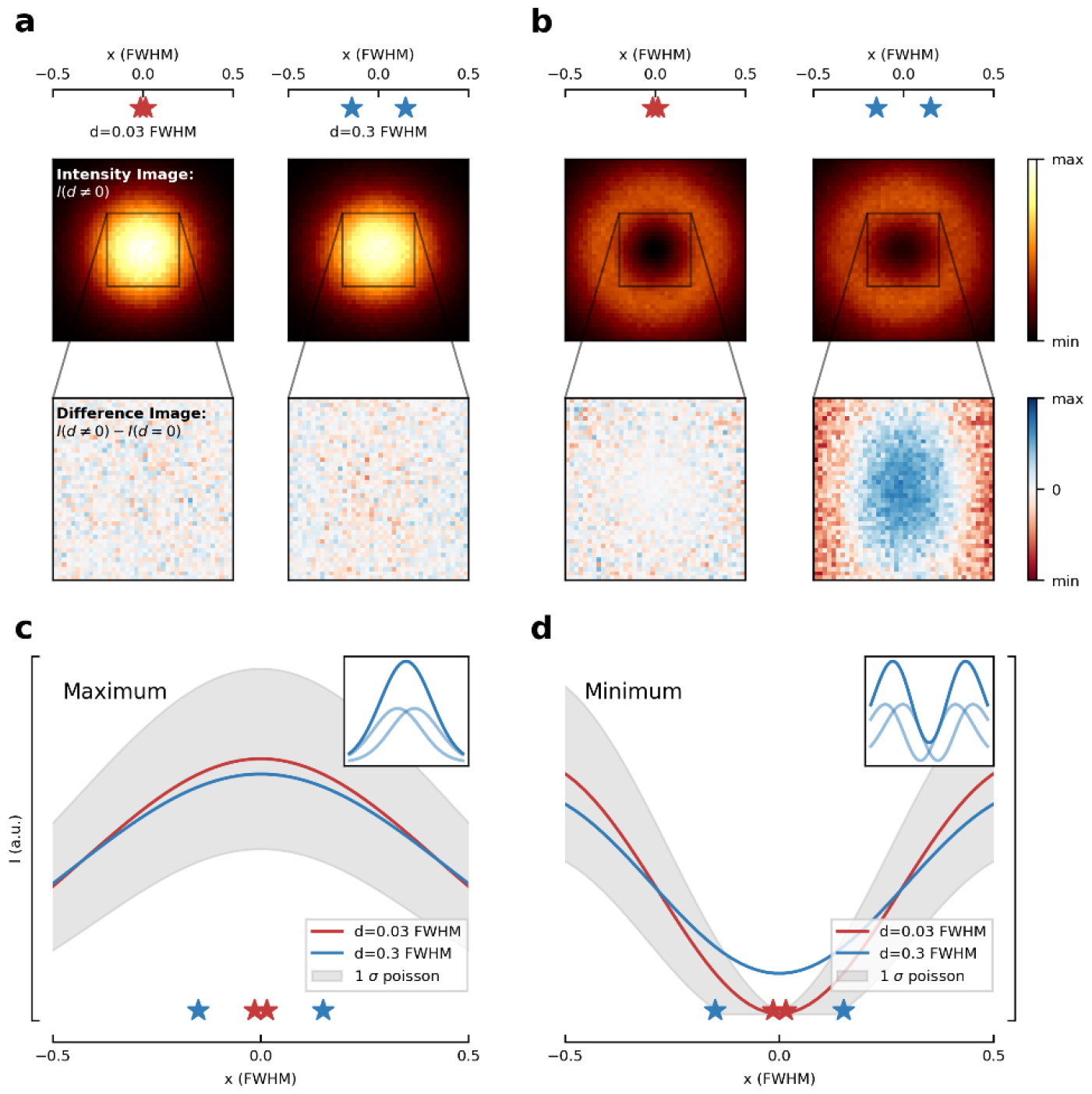
Utilizing a diffraction maximum vs. a minimum of light to resolve two inelastic point scatterers. **a:** When probed with a diffraction maximum of focused illumination light of certain Full-Width-Half-Maximum (FWHM), two closely spaced scatterers (illustrated as stars) cannot be resolved for separations below the diffraction limit of *d* ≈ 1 FWHM ≈ 280 nm. For separations below this limit, changing the positions of the scatterers only marginally alters the combined emission (note the similarity of the difference images for two separation values *d*_1_= 0.03 FWHM and *d*_2_ = 0.3 FWHM shown in the panel row below). For each image, *N* = 10^6^ detected photons are considered in the calculation. **b:** When probed with a minimum, the same disparity in *d* notably alters the joint signal; note the signal increase (blue shading) in the pertinent difference images. **c:** 1D Intensity profile of scattered light when illuminating the scatterers with a diffraction maximum. Changing *d* yields an intensity modulation of the joint signal that remains within the (Poisson) noise band of the mean signal for both *d*, i.e., the two sources cannot be resolved amid noise. **d:** When illuminating the same scatterers with a diffraction minimum, the modulation at the minimum of the resulting signal is outside the noise bands, allowing separation. Decreasing *d* results in a deeper minimum of the joint signal. The insets in panels c and d show the profile of the individual average intensity profiles scattered by each point scatterer as well as their joint signal.

In general, changing *d* between two inelastic point scatterers modulates the spatial distribution and amplitude of their joint image signal *I(d*). For overlapping diffraction maxima (Figure 1, c), the noise adds up and the modulation is noticeable only for about *d*≥*λ*/2. Fortunately, the modulation is easier to detect by contrasting the joint image signal with a zero-signal baseline. Harnessing this idea, we investigated the scanning of an excitation beam with a focal light field having a central intensity minimum (Figure 1, b). Since the resulting fluorescence signal is proportional to the illumination intensity, the signal of each individual fluorophore is minimal (zero) at the central node. Moreover, as the signal of inelastic scatterers adds up (incoherently), scanning over two scatterers with *d*<*λ* / 2 also yields a joint signal *I(d*) with a single minimum (Figure 1, d). In the absence of background this minimum is truly zero only for *d*= 0. Hence any deviation of the fluorescence minimum from zero is indicative of a finite *d*.

We first consider a sinusoidal excitation (fringe) pattern that is scanned over two fluorophores at distance *d* over a full period along the *x*-axis (Figure 1, d), for simplicity. The resulting joint signal is given by *I(d, ϕ*) = *a*_0_ + *a*_1_(d) cos(*ϕ* − *ϕ*_0_, with *a*_0_) denoting a constant offset and *a*_1_ being an amplitude varying with *d*. The parameter *ϕ*_0_ gives the phase difference of the joint signal with respect to the phase *ϕ* = 4π *x*/*λ* of the sinusoidal illumination pattern. *a*_0_ and *a*_1_ depend on a number of parameters, including the brightness of the fluorophores, their separation, background, and the (usually finite) initial fringe contrast of the illumination (see SI for detailed derivation). Furthermore, we denote the modulation visibility with *v*(d) = *a*_1_(d)/*a*_0_.

A finite *d* causes the sinusoidal fluorescence signals of the individual fluorophores to be spatially shifted with respect to each other, changing the visibility *v*(d) of the joint modulation. Interestingly, contrary to the case of separation by maxima, reducing *d* increases *v*(d), implying that measuring *d* with minima actually excels at small *d*. We recall that in the limiting case of *d*= 0 and zero background, the joint signal equals zero, meaning that even tiny *d* can be measured due to the intrinsically low noise at the minimum. By contrast, for overlapping maxima at *d* = *λ*/4, *v*(d) approaches zero and the separation is maximally challenged by noise. In any case, sampling *I(d, ϕ*) at three positions suffices to determine *a*_0_, *a*_1_ and *ϕ*_0_ and to resolve at *d* (see SI).

A lower bound on the precision with which the position of the two emitters can be estimated is provided by the Cramer-Rao Bound (CRB)^10^. The CRB is directly related to the Fisher Information (FI), which provides a measure on how much information can be inferred about the parameters of a model for *I(d, ϕ*), including the position of the fluorophores, given a statistical sample of their signal and a *I(d, ϕ*) model. We regard the inelastically scattered signal as an inhomogeneous Poisson process with an underlying mean specified by the convolution of the illumination pattern with the PSF of the imaging system.

For a Poisson process, the FI is proportional to the ratio of the model gradient (squared) ∇*I(d, ϕ*) and the absolute model value *I(d, ϕ*). This relationship maximizes the FI for scattered photons from the minimum and renders them more informative than those near the maximum. For two sources with separation *d*, the CRB as the precision σ of the distance estimate scales approximately as 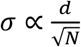 and 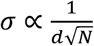 for the minimum and the maximum, respectively. Consequently, we obtain a constant relative error of the estimate of *d* for the minimum, but a diverging relative error for the maximum at small distances (Figure 2, a). This holds true only in the limit of an infinitesimally small scanning interval *L* → 0 and perfect contrast *v*_0_ = 1 of the illumination pattern. A larger (finite) scanning range *L* > 0 and an imperfect initial contrast *v*_0_<1 degrade the quality of the estimate (Figure 2, a-c). Yet, the estimate of *d* provided by a minimum is at least two orders of magnitude more precise compared to that afforded by a maximum (Figure 2, a). We note that this finding is irrespective of the shape of *I(d, ϕ*) since the decisive element is the minimum of the illumination beam, which, in first approximation, has a parabolic spatial intensity profile.

**Figure 2:**
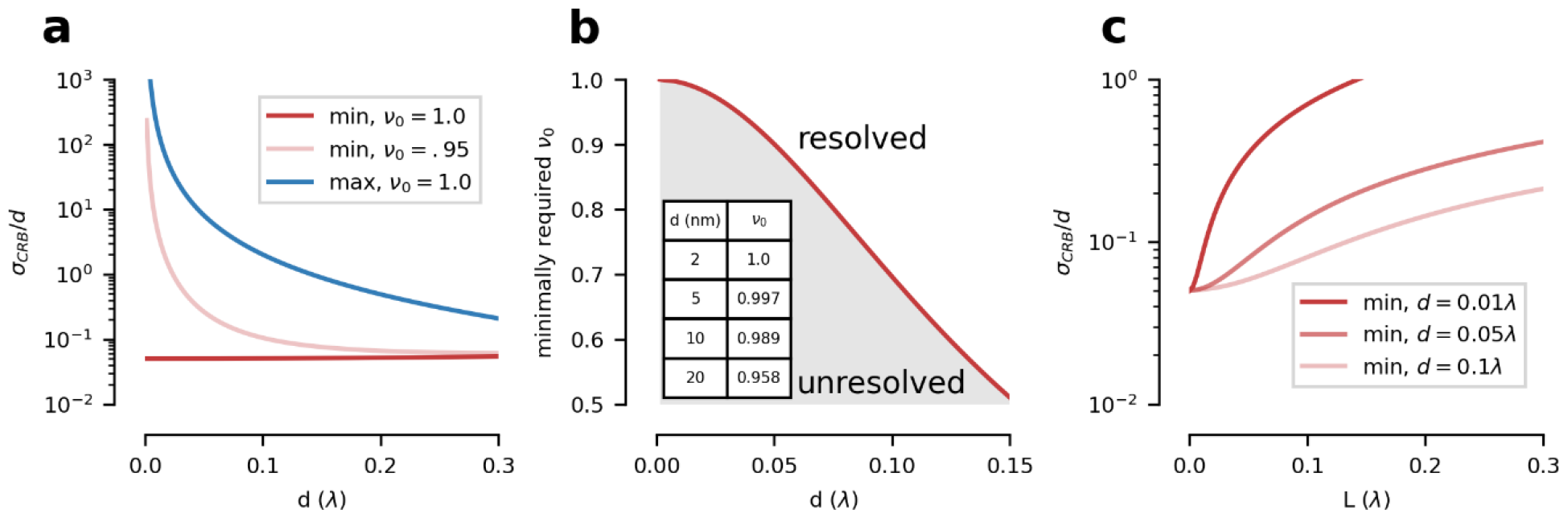
Theoretical localization precision of two point scatterers probed with a diffraction maximum (max) or a diffraction minimum (min) for N=100 detected photons. **a:** Cramér Rao-bound (CRB) divided by separation *d*, i.e., relative CRB (σ_*CRB*_ /*d*), for different initial visibilities *v*_0_. When probing with a minimum the relative CRB for the resolution remains constant, while it diverges for the maximum. Imperfect contrast of the minimum of the illumination light (*v*_0_=0.95) deteriorates the precision, yet the relative CRB is improved by roughly two orders of magnitude over its counterpart employing a maximum. **b:** Impact of the visibility *v*_0_ on the resolvable distance d. Measuring small distances requires a high contrast of the illumination pattern,i.e. a minimum with sufficient ‘depth’. Here, a successful distance measurement (‘resolved’) is required to exhibit a relative CRB<0.5. The inset table provides exemplary values of required minimum visibility to measure *d* (*λ* = 640 nm). **c:** Relative CRB with respect to scanning range *L* near the minimum of the combined signal, exemplified for various *d*. The precision improves with decreasing *L*, which implies that probing as close to the minimum of the joint signal improves the distance estimate.

The CRB does not guarantee the existence of an estimator that attains this precision bound. However, among a range of options (see SI), we found that a polynomial maximum likelihood estimator of the parameters *a*_0_, *a*_1_ and *ϕ*_0_ is sufficient to retrieve a *d* estimate from the photons near the minimum with a constant relative error. This advantage also remains amid non-negligible background and non-vanishing intensities at the illumination diffraction minima (see Figure 2, b and SI).

To experimentally find the minimal distance at which two inelastic scatterers can be distinguished, we placed two fluorophores (Atto647N molecules) at controlled distances 6 nm < *d*<90 nm, that is, at distances that are definitely considered not resolvable due to Rayleigh’s limit. This spatial arrangement was realized by attaching the fluorophores to a DNA origami structure, a so-called nanoruler^11^. The DNA structure served as a scaffold for attaching the fluorophores at a given *d*. Next, we employed a line-shaped interference pattern in the focal plane of a scanning confocal microscope. The pattern was obtained by illuminating opposing halves of the entrance pupil of its 1.4 numerical aperture oil immersion lens with two interfering excitation beams (Figure 3, a). Initially used for MINFLUX^12^, this setup was detailed previously. In brief, the line-shaped focal interference pattern was oriented either in the x- or the y-direction and could be quickly interchanged by changing the x-and y-orientation of the two beams using an electro-optic device (Figure 3, b). Destructive interference at the focal point provided a central line-shaped diffraction minimum, whereas constructive interference a maximum. Changing the phase difference linearly between the beams allowed to scan the pattern over the fluorophores (along the x- or the y-axis, Figure 3, c) so that we could readily compare the separation results provided by the minimum with that by the maximum.

**Figure 3:**
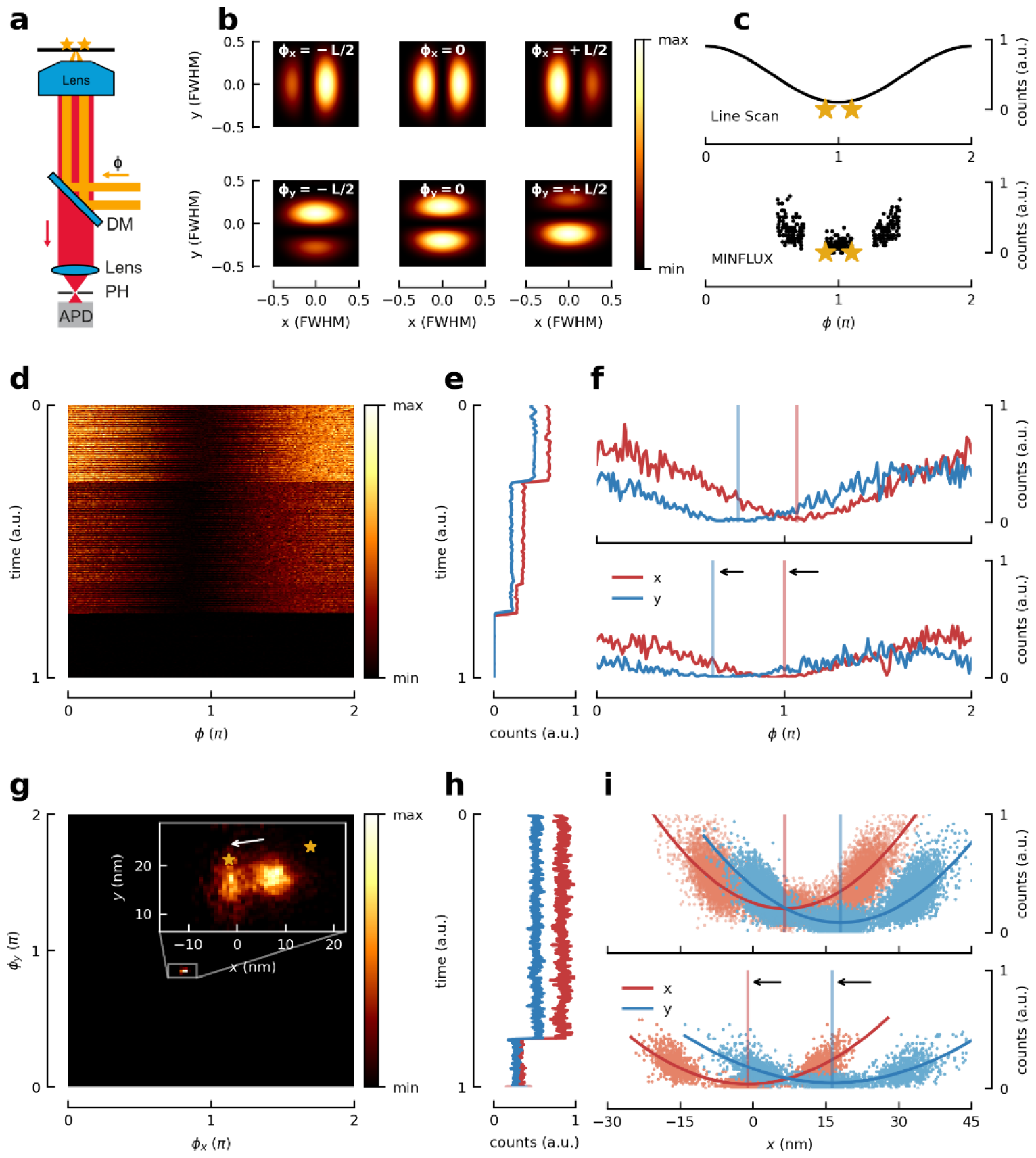
Measurement of distances between two simultaneously emitting fluorescence molecules by (x,y-) scanning with an illumination intensity minimum. **a:** Scanning fluorescence microscope with photon-counting detection (APD) of fluorescence passing the dichroic mirror (DM) and a confocal pinhole (PH). The interference of two beams with adjustable phase difference *ϕ* entering the pupil of the objective lens creates an illumination intensity pattern in the focal plane featuring x- or y-oriented line-shaped diffraction minima and maxima (MINFLUX setup). Two fluorophores sketched as stars **b:** Changing *ϕ* scans the line-shaped minima in the y-and x-direction. **c**: Top: Line-scan principle: linear ramp of *ϕ* over 2π shifts the minimum across the scatterers, producing a sinusoidal line profile of fluorescence (or scattered signal). Bottom: The continuous line-scan is adequately replaced by probing three points near the scatterers with the minimum (MINFLUX recording). **d**: Normalized counts measured during repeated line-scans. The absolute number of counts decreases over time in a stepwise manner as individual fluorophores bleach. Repeated ramping of *ϕ* over 2π in x- and y direction across the scatterers yields a line-scan stack. **e:** Averaged counts per line, normalized over the whole stack. Two bleaching steps are clearly visible, marking transitions from two molecules, to one, to zero (background). **f:** Exemplary lines from two and a single molecule show the sinusoidal profile of the fluorescence counts and the spatial centre-of-mass shift after the first bleaching step. Each line allows to extract information based on the photons just near the minimum, the maximum, or from the entire line. **g:** Heat map of localizations of the fluorescence center-of-mass for two fluorophores at 20 nm distance. Inset: Two clusters of localizations are visible showing the center-of-mass shift after one fluorophore bleached. **h:** Averaged counts per 3-points MINFLUX measurement for x and y axis normalized over the whole measurement. The bleaching steps and fluctuations in fluorophore brightness are clearly visible. **i:** Normalized counts for each segment. A second order polynomial fit shows a change of the shape of the parabola after the bleaching step as well as a shift of the position of the minimum. Measurement of *d* relies only on photons from the first (two molecule) segment; the bleaching steps are considered just for an independent control.

These line-scans were consecutively repeated for both the x- and the y-axis until both molecules bleached; the stack of all x- or y-line-scans constituted an x- or y-trace, respectively (Figure 3, d). Thus, each trace allowed us to consider all photons or just those originating near the diffraction minimum. For both cases we determined a projection of the distance between the fluorophores to each axis and calculated *d*. We also recorded the sudden bleaching of individual fluorophores during the measurement (Figure 3, e). The ensuing sudden shift of the center of mass of the signal allowed us to extract *d* as well (Figure 3, f-i). Pioneered in camera-based single molecule localization^13^, this ON/OFF-based separation provided an independent control of our *d* measurement.

Our experiments show that separating by scanning the entire interference pattern over the sample becomes inaccurate for *d* < 30 nm, whereas selecting photons from a region just near the minimum of a stacked line scan (trace) resolves the fluorophores down to *d* of 10 nm. Moreover, the results nicely agree with those obtained by the bleaching control (Extended Data Figure 1). In accordance with theory, this finding demonstrates that the photons registered from the minimum excel at resolving fluorophores at small *d*. Perhaps more strikingly, using additional photons not only consumes time and the limited fluorescence budget, but also compromises resolution.

Consequently, we next probed the two fluorophores just with an excitation minimum. Like in previous MINFLUX recordings, we sampled three positions at 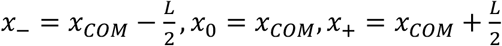 in each axis whereby the size of *L* was iteratively reduced. The center-of-mass (*x*_*co*m_) of the two fluorophores was subsequently estimated using the fluorescence minimum. To estimate *d*, a number of such triples of positions and counts is combined into a bin of a certain number of photons. This procedure allows reaching single-digit nanometer accuracy for the *d* estimate with about 5000 photons (Figure 4). Besides, it paves the way to resolving in dynamic settings, where distance and location of two fluorophores have to be determined continually, i.e., without ON/OFF interruption.

**Figure 4:**
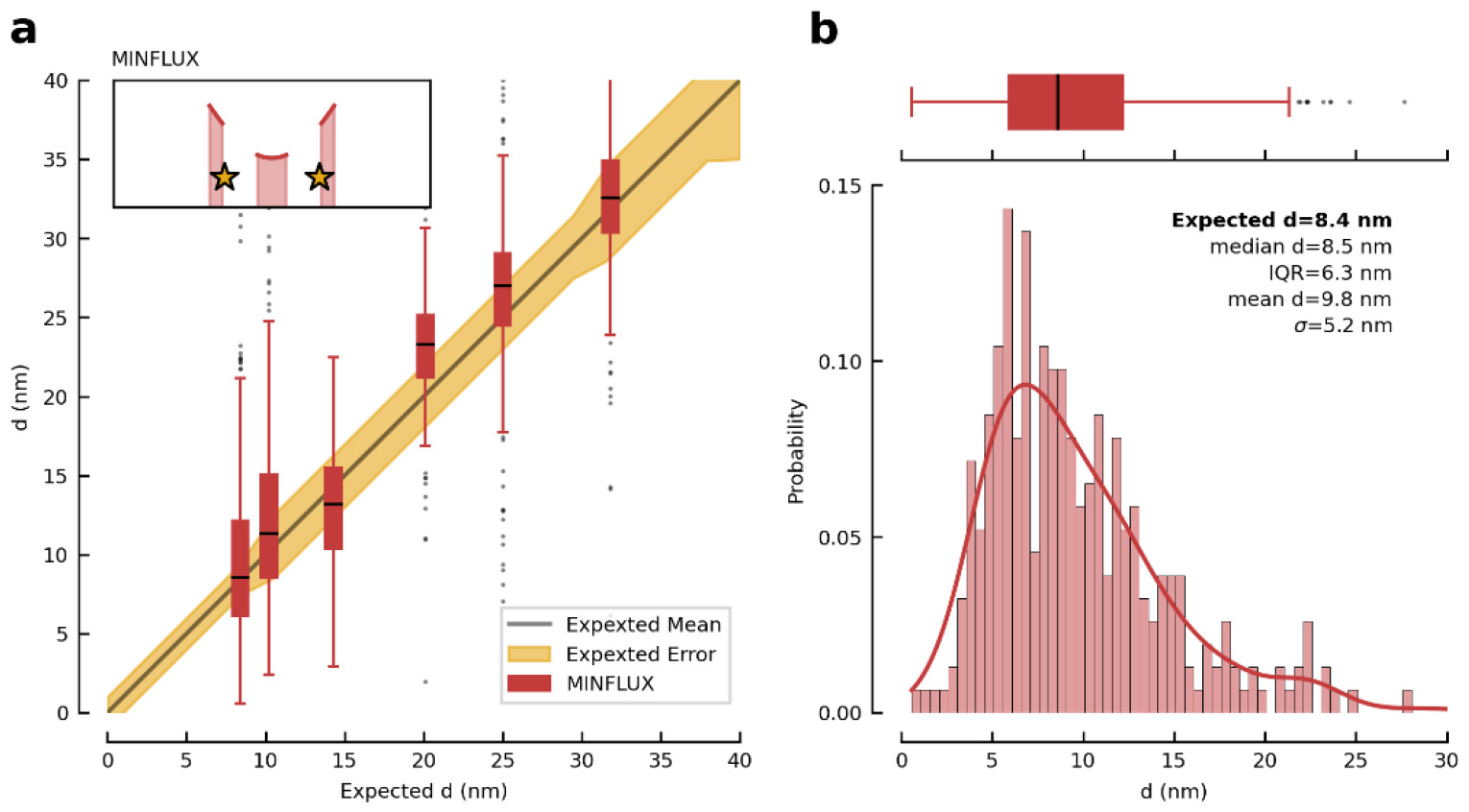
Resolving two constantly emitting fluorophores at distances down to 8 nm. **a:** Boxplot of the measured distances *d* over the expected distance (black line as specified by the manufacturer of the DNA nanoruler scaffold). Recording the photons originating near the illumination minimum (see inset) suffice to separate simultaneously emitting fluorophores down to *d* = 8.4 nm within the uncertainty specified by the manufacturer (yellow shaded area). The box extends from the lower to upper quartile values of the data. Error bars mark the interval from the median (black line within box) to the last point within a 1.5 interquartile range (IQR). Outliers are shown as black dots. Independent measurements per box: 8.4 nm: 307; 10.2 nm: 322; 14.3 nm: 218; 20.1 nm: 272; 25 nm: 231; 31.8 nm: 283. **b**: Histogram of the distances obtained for the 8.4 nm fluorophore separation overlaid with the scaled probability density function (red line). The corresponding box plot from a is shown above.

To demonstrate tracking of two moving emitting fluorophores, we translated the sample stage on a defined trajectory. As they were attached to a nanoruler, each of the fluorophores produced a slightly shifted copy of the trajectory of their counterpart. The time-resolved distance between the fluorophores can be extracted by tracking their center-of-mass and evaluating the signal for given points in time. Distance estimates for fluorophores at *d* = 30 nm (moving along a circle with 30 nm of diameter) and at *d* = 15 nm (moving on a higher-order Lissajous trajectory with 15 nm amplitude) are readily obtained (Figure 5, Supplementary Video). These results, which are clearly not possible with popular camera-based tracking, demonstrate time-resolved distance measurement of spectrally identical and constantly emitting fluorophores far below the diffraction barrier. In other words, for establishing the nanometer scale distance as a function of time, ON/OFF switching or separation by excitation/emission wavelengths are not needed. This finding greatly relaxes the requirements on the fluorophores, since stable non-switching and non-blinking fluorophores can be used. Stable fluorophores generally bring about higher photon emission rates and much larger photon emission budgets. Moreover, individual fluorophores vary much less in emission which improves the precision of the distance estimates.

**Figure 5:**
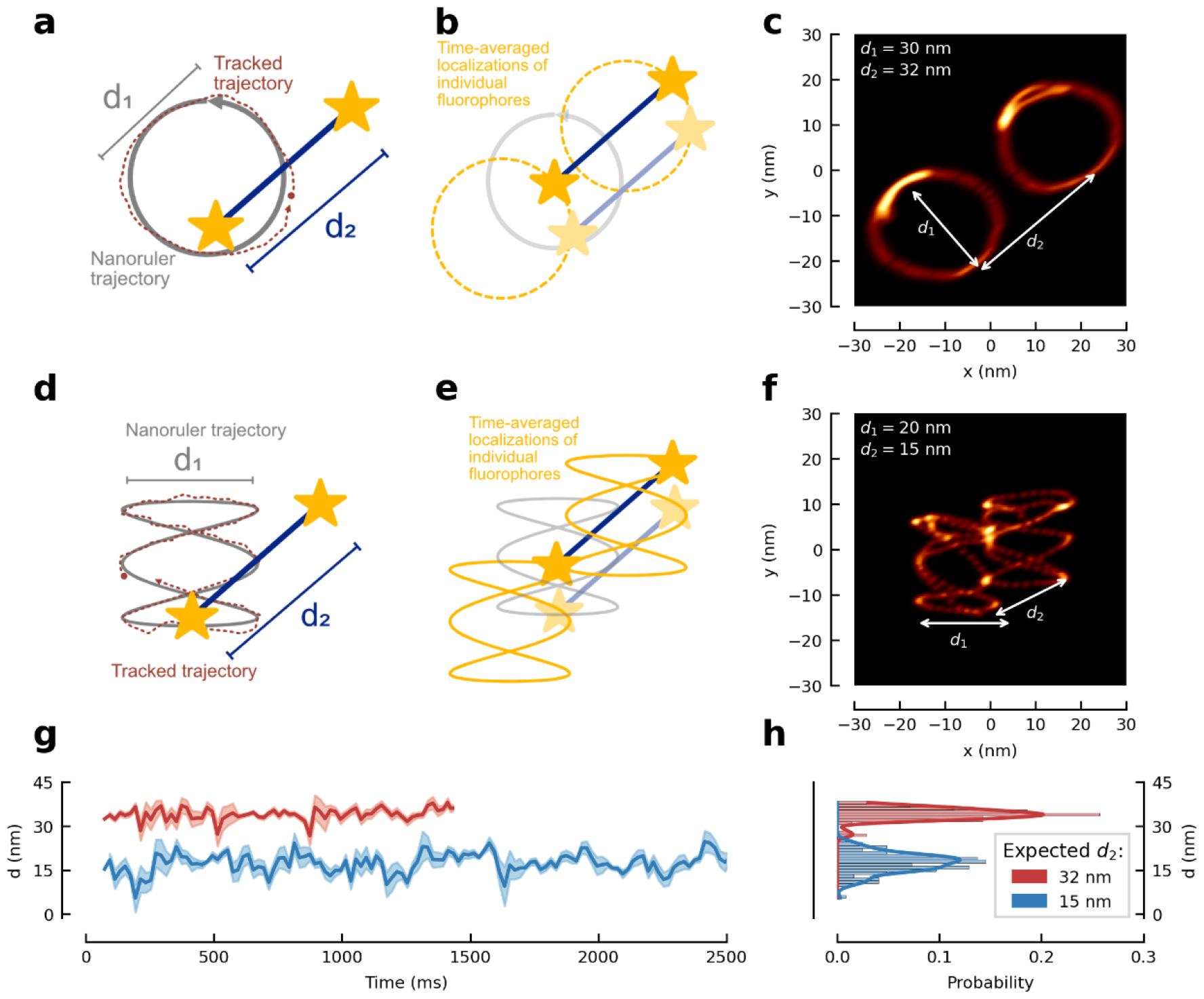
Uninterrupted resolution and localization of two moving fluorophores. **a**: Schematic of the measurement: A nanoruler with two fluorophores is moved on a pre-defined trajectory (circle) using the microscope stage. The centre-of-mass of the two fluorophores is tracked with an illumination intensity minimum (dotted red line). Distances *d*_2_ are estimated in a bootstrapping approach along the obtained traces. **b:** The subsequent positions of the fluorophores are calculated relative to the current centre-of-mass coordinate. This results in the trajectories of the individual fluorophores (two yellow circles). **c:** Exemplary trace of a *d*_2_ = 32 nm nanoruler moving along a circle with *d*_1_ = 30 nm diameter. Time-averaged image of the two fluorophores results in two displaced circles. **d**: Nanoruler with *d*_2_ = 15 nm moving along a Lissajous-figure with amplitudes 15 nm and 20 nm in x and y direction, respectively. **e:** The trajectories of the individual fluorophores result in shifted copies of the centre-of-mass movement. **f:** Exemplary trace of a *d*_2_ = 15 nm nanoruler moving along the Lissajous figure. Time-averaged image of the two fluorophores results in two displaced copies of the Lissajous figure. **g**: Time series of measured distances (thick line) for 50 ms time-bins. Error bands correspond to a moving standard deviation (200 ms window). For each bin, we select the respective tuples according to their time-stamp and estimate the distance and orientation of the fluorophores. **h**: Histogram of measured distances (no averaging) yields clearly distinguishable populations for nanorulers of *d* = 32 nm (red) and *d* = 15 nm (blue). The histograms are overlaid with the scaled probability density function.

A remaining question is whether the potential of the diffraction minima can be extended to identifying the position of three or more identical scatterers located in a given geometric constellation and distance within regions ≪ *λ*/2. To address this question, we simulated various constellations of equally bright fluorophores (Figure 6). The arrangements are parametrized by a single scaling parameter *d*. For a linear arrangement of scatterers, *d* corresponds to a nearest-neighbor distance or it is the diameter of the circle circumscribing m scatterers forming a regular polygon. We found that the advantageous scaling of the relative error σ/*d* = *RMSE*(*d*)/*d* remains valid for more than two scatterers. Conceptually, a smaller distance parameter *d* allows to discern a larger number of scatterers. In practice, the attainable precision is lower-bounded by experimental imperfections such as finite background, variation of individual source brightness, and reduced initial contrast of the illumination pattern, i.e., finite ‘depth’ of the intensity minimum. Yet, our numerical data shows that for *d* ≪ *λ* / 2 and commonly encountered SNR, the position of individual inelastic scatterers can be identified for various geometrical arrangements. For scatterers arranged along a line at a given recurring distance *d*, the relative error increases with *d* as the ensemble substantially extends into areas of larger intensity outside the probing minimum. Interestingly, not only linear but also polygonial or grid-like arrangements of a known number of identical scatterers allow the identification of their mutual distance with a given number of detected photons.

**Figure 6:**
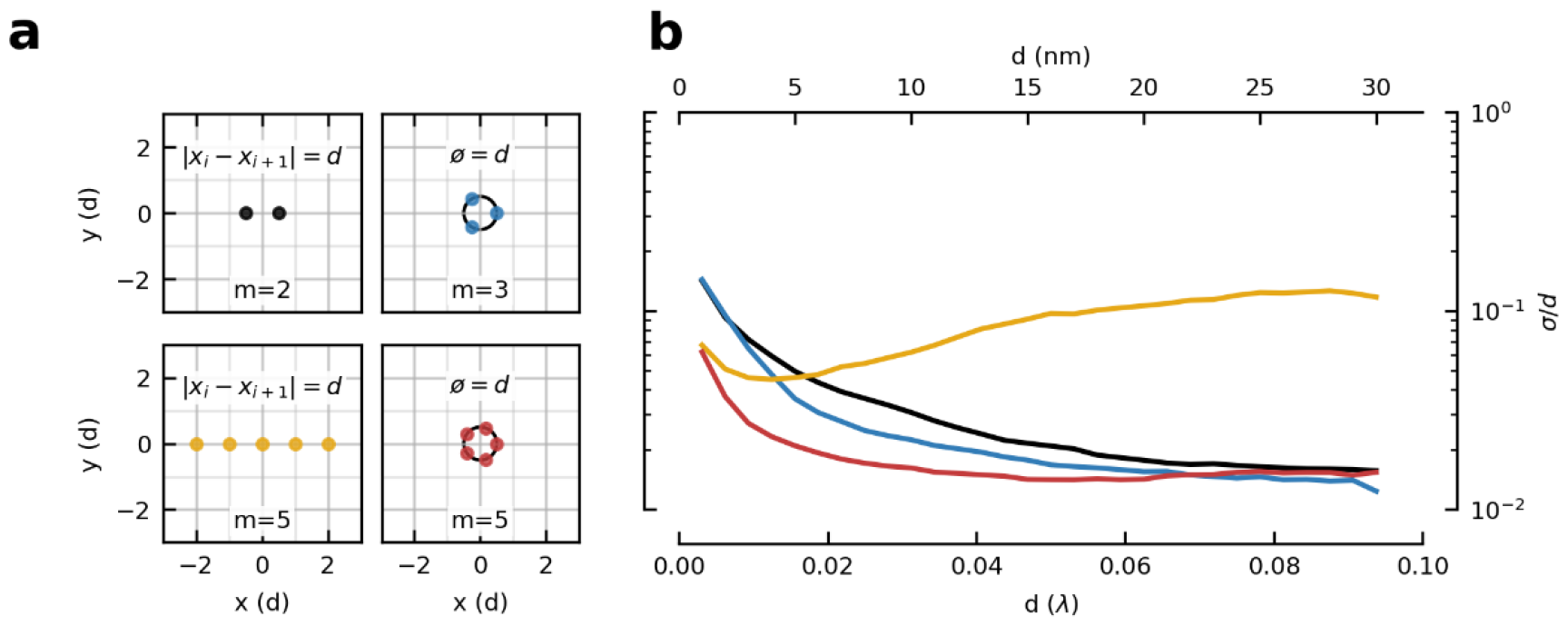
Sub-diffraction localization of multiple simultaneously scattering point sources; numerical simulation. **a:** Spatial arrangement of up to m=5 point scatterers (fluorophores): lines and regular polygon. The scaling parameter d denotes the nearest-neighbour distance between point sources on a line or, in case of the polygon, the diameter of the circle circumscribing all sources. **b**). Numerical simulation of the relative distance error σ/*d* = *RMSE*(⟨*d*⟩)/*d* for the arrangements shown in a (same color coding). Note that at distances *d*<0.018*λ*, five scatters in a line can be resolved more precisely than two, indicating that the resolution favourably depends on the number of scatterers as long as the ensemble does not significantly extend beyond the minimum (*d* > 0.02 *λ*) of the probing intensity beam (compare black and yellow line). In case of the polygon, adding a scatterer increases the emitter density i.e., does not change the spatial extent of the ensemble, and hence improves the accuracy of the *d* estimate at small *d* (compare red with blue line). Again, this implies that over small distances *d*, for example *d*<20 nm, localization and measurement of distance between a larger number of scatterers is advantageous over a smaller number. The numerical distance values are obtained for *N* = 500 detected photons, a realistic probing range of *L* = 30 nm, an average background emission equivalent to *β* = 0.1 emitting molecules in the background and *λ* = 640 nm.

In a nutshell, employing diffraction minima instead of maxima allows in principle diffraction-unlimited separation of two spectrally identical incoherent point scatterers – even at finite photon numbers. This finding holds also for an arrangement of a known number of scatterers that are confined to sub-diffraction regions at distances *d* ≪ *λ* / 2, provided the geometrical form of the arrangement is by and large known. This finding has great potential for the optical investigation of dynamic (re)arrangements of proteins and other molecules with conventional optics at the nanoscale.

Moreover, our study highlights that, owing to the wide acceptance of the Rayleigh criterion, the ability of resolving individual scatterers with freely propagating light waves has been profoundly underestimated. We think that this is partially due to the fact that when employing diffraction maxima, resolution becomes increasingly harder with decreasing *d*, rendering the vision of extracting dimensions *d* ≪ *λ* / 2 with propagating (light) waves unrealistic. In stark contrast, our work shows that measuring with a diffraction minimum instantly opens up a measurement window in the *d* ≪ *λ* / 2 range resolving distances at 1/80 of the wavelength.

Besides the lower noise level at the minimum, a major component to the success of our method is the robustness of the minimum with respect to aberrations. The spatial intensity profile of the minimum is approximately parabolic and is sampled only in the vicinity of the minimum, which makes it less prone to distortions. In comparison, popular deconvolution approaches employing diffraction maxima require detailed knowledge of the (aberrated) PSF of the system and need to identify these details amid the shot-noise at the maximum. Since there is no fundamental difference between fluorescence and other types of inelastic optical scattering, e.g. Raman scattering, given similar signal and background conditions, our results should also be transferrable to these important fields of optics.

Last but not least, it has not escaped our attention that our principle of separating with minima should also be applicable to elastic scatterers at small distances, where distance-dependent phase differences between the light scattered by different sources should be negligible. Equally intriguing is the fact that by avoiding state transitions of the investigated material for separation, our approach of identifying the position of point scatterers with a diffraction minimum should be applicable to any type of point scatterers of any type of propagating wave, thus opening up superresolution to many other wave-based imaging modalities.

## Acknowledgements

T.A.H. is part of the Max Planck School of Photonics supported by BMBF, Max Planck Society, and Fraunhofer Society. The authors wish to thank their colleague S. J. Sahl for helpful discussions and critical reading of the manuscript.

## Author contributions

T.A.H. performed the theoretical analysis, designed and carried out the simulation and evaluated the experimental data. J.O.W. performed the experimental investigation, supported by T.A.H, and built the utilized setup. S.W.H. outlined the idea of using a minimum to separate identical emitters and steered the investigation. All authors discussed the results and their implications during the course of the project. The manuscript was written by T.A.H. and S.W.H.

## Methods

### Experiments

#### Experimental Setup

The measurements were performed on a previously described setup^1^ using the interference between two horizontally or vertically displaced laser beams generating an excitation beam profile with a line-shaped minimum oriented in either x- or y-direction in the focal plane, respectively. This minimum can be moved in the focal plane along the axis of beam separation by electro-optically changing the phase difference between the two beams.

#### Line-Scan Measurements

After a coarse pre-localization step of the center-of-mass of the linear nanoruler featuring a fluorophore at each end, the illumination pattern is alternately scanned over the nanoruler in x- and y direction. When changing the phase difference linearly, and thus linearly moving the line-shaped minimum across the focal region, the emitter is subjected to a sin^2^-intensity modulation referred to as ‘line-scan’. A full period (320 nm) of the harmonic signal is recorded, discretized by 160 pixels and a dwell time per pixel of 500 µs. The procedure is repeated 300 times (total scan time of 48 s) at the same position until both molecules bleached. The stack of all one-dimensional (1D) line-scans of x- and y-axes forms the x-and y-trace, respectively. Due to the short acquisition time, the measurement was not particularly stabilized against mechanical or thermal drift. The laser power during the phase scans was set to 20 µW, which resulted in an average count rate around 70 kHz.

#### Nanoruler selection

To record individual nanorulers, a 10×10 µm^2^ field of view was scanned with an average continuous-wave laser power of 2 µW. After smoothing the image with a Gaussian kernel, the spots were iteratively selected by fitting a region around the brightest pixel with a Gaussian, subtracting the fit and repeating the procedure until 10 spots were found. Those spots were addressed by the galvanometer-scanner to perform the linear phase scans along the x- or y-axis.

#### Line-Scan Analysis

Line-scan traces are composed of 1D line-scans in the x- and y-axis, which we processed independently to obtain a distance estimate in each axis and calculate the final distance as their norm. In the following, we describe the routine for an individual axis. Traces are segmented with a change point detection routine, irrespective of the count rate. (Python package *ruptures*^2^, piece-wise constant model and a penalized segmentation routine (PELT) to detect a previously unknown number of change points). The trace is split at the detected change points, and each segment truncated five lines before and after the bleaching step. Ideally, the trace has three segments. More segments are defined depending on the number of molecules, bleaching steps or marked changes in brightness.

The traces are pre-selected by eye in order to discard obviously faulty measurements, e.g., as in the case of a bright cluster of molecules or a lack of bleaching steps (only one molecule was present or the two molecules did not bleach during the measurement). Next, we automatically filter for traces with initially two molecules, two bleaching steps and eventually zero molecules (background). Additional automatic filtering is performed on the segments to restrict the standard deviation of the average counts to less than 10 times the expected Poisson noise of the mean value of the line (to filter for short blinking events). The ratio of counts between the two-molecule and the one-molecule segment is allowed to deviate by ±50%. The automatic filter routine has a success rate of 96% on the data that has been pre-selected by eye.

Once the filtering is complete, we transit to the evaluation of the line scans. Since the selected traces have both a two-molecule and a single molecule trace, we perform a local calibration for each phase scan by estimating the local background from the corresponding background segment of the trace and the quality of the intensity minimum (initial fringe contrast) from the single molecule segment. This estimate serves as calibration for the following evaluation of the two-molecule segment. The two-molecule segment is evaluated in a bootstrapping approach to obtain an estimate of the reduced two-molecule visibility by fitting different models to the data, e.g. a polynomial ansatz near the minimum or a harmonic model considering all photons of a line (see SI for all available methods we tested). The locally calibrated initial visibility and the reduced two-molecule visibility allow calculating the projection of the distance in the respective axis. All analysis routines were implemented in python.

#### MINFLUX-like measurements

We leverage an iterative MINFLUX-like approach, i.e scanning with intensity values around the zero point, in order to sample the two emitters only in a region around the minimum of the illumination pattern. In contrast to a full line-scan, this drastically reduces the number of photons collected, increases the speed of the measurement and ensures the collection of the most informative photons. The actual MINFLUX routine consists of 4 steps with different *L*-values, laser powers and number of repeats. In each step the emitter is repeatedly localized by probing at positions 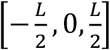 around the current position estimate *x*_*est*_ in first x- and then y-direction. Each exposure lasts 158 µs which (together with wait times for addressing new positions) results in a total time of around 1 ms for a complete xy-localization. Based on the recorded photons the position estimate is refined after each localization according to

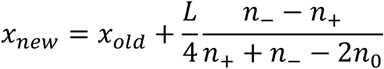

with [*n* −, *n, n*] being the photons recorded at 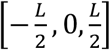 repectively. Localizations are considered as failed if either less than five photons are collected or if 2(*n*_+_ + *n*_−_ − 2*n*_0_)<|*n*_−_ − *n*_+_| which limits the allowed corrections to *L*/2. While the position of the center-of-mass is estimated and targeted anew after each measurement triple, the probing range *L* is decreased only when progressing to the next step. The first steps serve to quickly localize the center-of-mass of the system (position of the minimum) and “zoom-in” with the intensity minimum, only photons of the last iteration are used to estimate the separation of the source. In total 5000 localizations are performed resulting in a total measurement time of around 5 s. An overview over the *L*-values, laser power and repeats is given in table S1.

#### Analysis of MINFLUX recordings

Measured traces consist of tuples denoting probing position of illumination pattern, time and measured counts. Traces are segmented via a change point detection algorithm (python, ruptures). Segments are classified as background, single molecule traces or two molecule traces depending on the average count rate of the segment and subsequent bleaching steps before or after the segment. Segments are further treated in a bootstrapping approach to create chunks with a certain photon number, number of measurement tuples or time-bins. The mode is chosen depending on the experiment. In the case of static distance measurement a photon-binning is appropriate, while dynamic settings require a time-binning. Each chunk is fitted with various models, e.g. a polynomial estimator near the intensity minimum. Failed fits (those that did not converge) are excluded from further analysis.

To calibrate, we calculate the average background counts from the segments containing only background for each measurement batch (e.g. size of nanorulers) and in the respective axes. This allows calculation of the background-corrected single molecule visibilities *v*_0_ from the fit parameters of the segments containing only single-molecule data. To obtain a calibration value for each batch (size) and in each axis (x,y), we take a *X*^2^-weighted median of the visibilities within each batch and axis. Note that the calibration could also be performed in a separate measurement and that it characterizes a property of the instrument. Last, we calculate the reduced visibilities on the two-molecule segments from the obtained fit parameters on the two-molecule segments. Calibrated by the background corrected initial visibilities of the batches and in each axis, we calculate the distance projections as well as the norm for each chunk. These norms are the final distance estimates of a nanoruler with respect to the chunk id (e.g. time) and allow a time-resolved co-localization. All analysis routines were implemented in python.

#### Control Method: Distance measurement via bleaching steps

Traces that contain at least one bleaching step offer an elegant way to run a control method for the distance estimate on the same data as the estimate with photons from the two molecule segment only. Since it is unlikely that both molecules bleach at the same time, many traces contain a two-molecule segment as well as a single molecule segment. While the minimum of a line-scan is located at the center-of-mass of the system when both molecules emit, it jumps to the position of the remaining molecule once the other one has bleached. Hence, observing this center-of-mass shift offers an independent control for the distance estimate. By employing a polynomial model that is fitted to the minimum, we extract the position of the minimum at any given time, average over the center-of-mass positions of the two-molecule trace and do the same for the single molecule trace after the first bleaching step. The molecule distance can be estimated as twice the lateral shift of the minimum.

#### Stage Movement for Dynamic Measurements

To create movements of the nanoruler and hence of the fluorophore, the stage is moved on a Lissajous curve. Therefore, the position of the stage is updated every 10 ms by adding the derivative of the desired trajectory 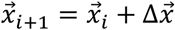. Assuming a Lissajous curve of the form

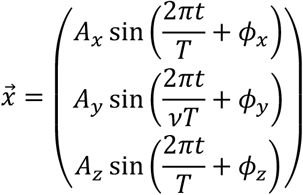

with amplitudes *A*_*x,y,z*_ a period of length *T* and potential phase shifts *ϕ*_*x,y,z*_ in each direction. The parameter *v* controls the ratio between the modulation frequencies and defines the lissajous shape. Then the derivative added to the current position is given by

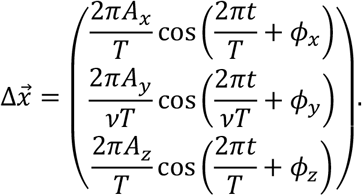

The Lissajous parameters used in this work are summarized in table S2.

#### Numerical simulations of multi-emitter ensembles

Results for the precision of the *d* estimate for more than two emitters were obtained via a Monte-Carlo simulation. We arranged emitters along a line, a regular polygon or a grid-like structure. We then simulated an iterative MINFLUX sequence resulting in a trace of tuples with positions of the illumination pattern and detected photon counts. This trace is chunked in a bootstrapping approach, conditioned to a maximum number of photons *N*. That is, we combined as many tuples as necessary until the sum of all collected photons reached the required value. The distance parameter *d* is then estimated on these chunks, and the Root-Mean-Square-Error (RMSE) of the estimates is calculated. The relative error is given by the fraction of the RMSE and the ground truth value of the parameter *d*.

#### Sample Preparation

##### Cleaning Process of Coverslip

To clean the cover slips (170 μm, No 1.5H, Paul Marienfeld GmbH & Co. KG, Lauda-Königshofen, Germany) measuring 15-18 mm in width, the following procedure was followed. Initially, they were gently wiped using a lint-free cloth sprayed with acetone. Subsequently, another cloth sprayed with Isopropanol was used to wipe them. After rinsing the cover slips with Isopropanol, they were dried using a nitrogen flow. Finally, a plasma cleaning process (using oxygen) was carried out for 5 minutes at 200 W.

##### Construction of Flow Chamber

To facilitate the easy exchange of incubation solutions during preparation and buffer changes between experiments, flow chambers were created. This involved attaching two narrow strips of double-sided tape (Scotch Double Sided Tape, 3M, Saint-Paul, USA) to an microscope slide, forming a channel approximately 5 mm wide. A cleaned coverslip was then placed on top of the tape and secured by applying gentle pressure.

##### Preparation of Nanorulers

Preparation of the DNA structures (Nanorulers, GattaQuant GmbH, München, Germany) was carried out with the following routine (see supplementary Table S3 for manufacturers of the substances):

1. Clean Coverslip
2. Prepare flow channel
3. Pipette the following through the flow channel:
  a. 200 μL PBS (flushing the flow channel)
  b. 15 μL BSA-Biotin (1:2 in PBS), wait two minutes
  c. 200 μL PBS (flushing the flow channel)
  d. 15 μL Strepatvidin (1:2 in PBS), wait two minutes
  e. 200 μL PBS (flushing the flow channel)
  f. 15 μL GattaQuant Rulers (diluted 1:50 or 1:100 in PBS), wait two minutes
  g. 600 μL PBS (flushing the flow channel)
  h. 15 μL Buffer
  i. Seal edges with epoxy
  j. Store in fridge

The buffer consists of an imaging buffer (IB), Methylviologen (MV, 75365-73-0, Sigma Aldrich, St. Louis, USA) and an enzyme system. It is prepared as follows: mix 100 μL IB, 1 μL MV and 1 μL enzyme system. Reagents for the buffer:

- Imaging Buffer:
  - PBS (basis)
  - 10% Glucose (wrt mass)
  - 2 mM Trolox (wrt final concentration in IB)
  - 0.2 M MV
  - enzyme system: 10 mg Pyranase Oxidase + 170 μL PBS + 80 μL Catalase

Additionally, pre-mouted samples of nanorulers were purchased directly from GattaQuant.

## Supplementary Tables

**Table S1.** Overview over the *L*-values, laser power and repeats used in the different MINFLUX steps.

| Step | $L$ | Laser power ($\mu\text{W}$) | repeats |
| --- | --- | --- | --- |
| 1 | 200 | 4 | 3 |
| 2 | 120 | 10 | 4 |
| 3 | 60 | 40 | 3 |
| 4 | 30 | 160 | 4990 |

**Table S2.**
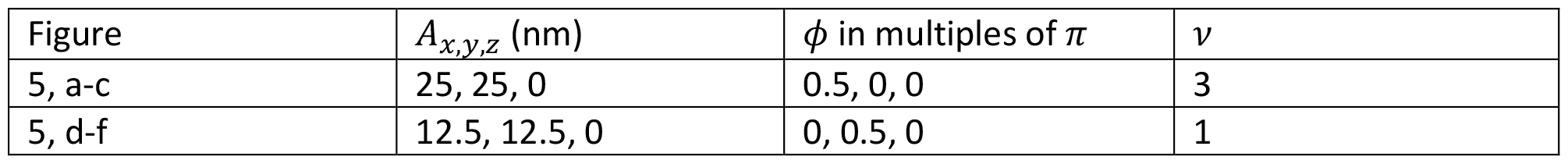
Overview over the Lissajous parameters used in this work.

| Figure | $A_{x,y,z}$ (nm) | $\phi$ in multiples of $\pi$ | $\nu$ |
| --- | --- | --- | --- |
| 5, a-c | 25, 25, 0 | 0.5, 0, 0 | 3 |
| 5, d-f | 12.5, 12.5, 0 | 0, 0.5, 0 | 1 |

**Table S3.**
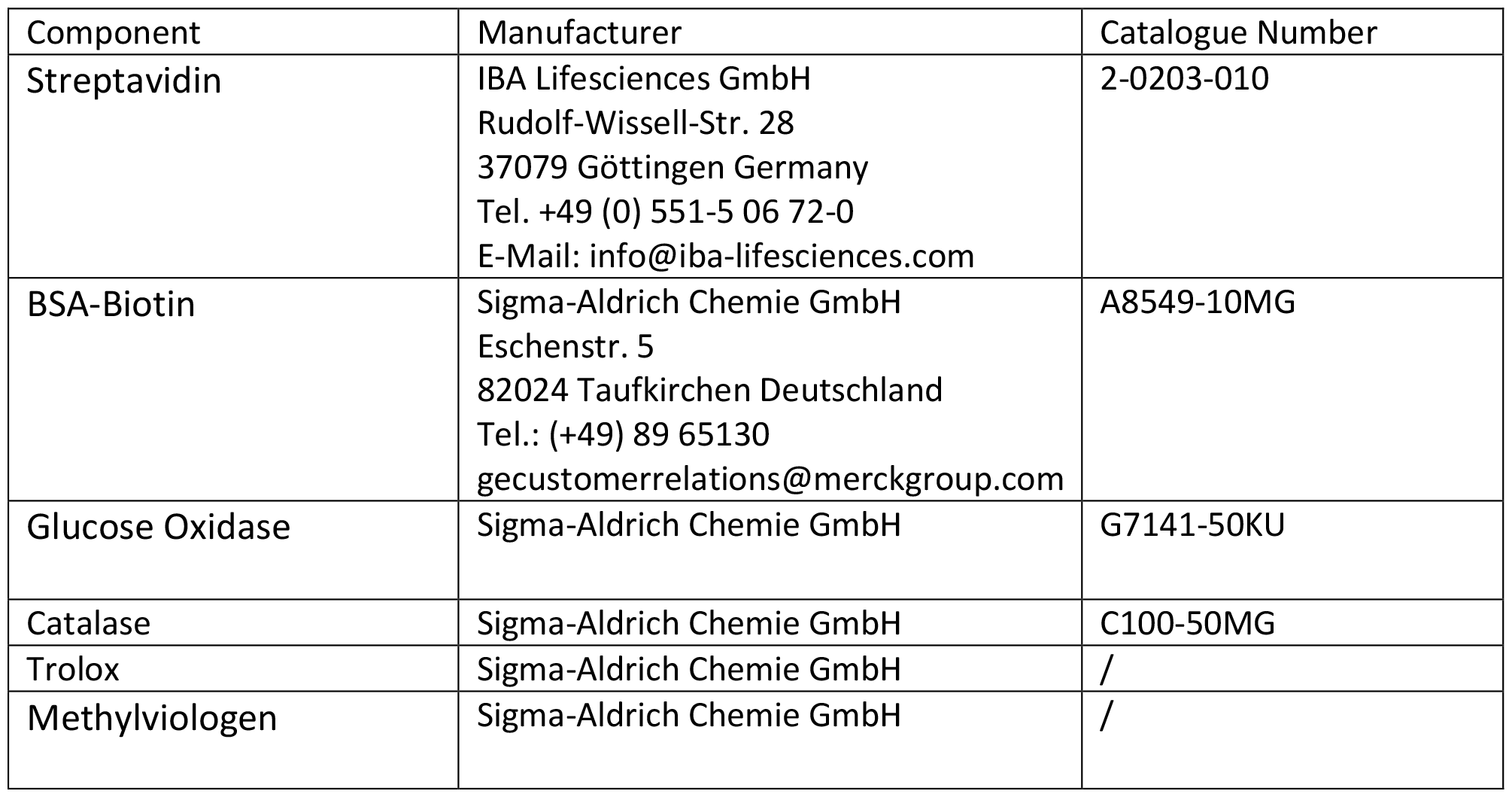
List of substance manufacturers.

---

**Extended Data Figure 1:**
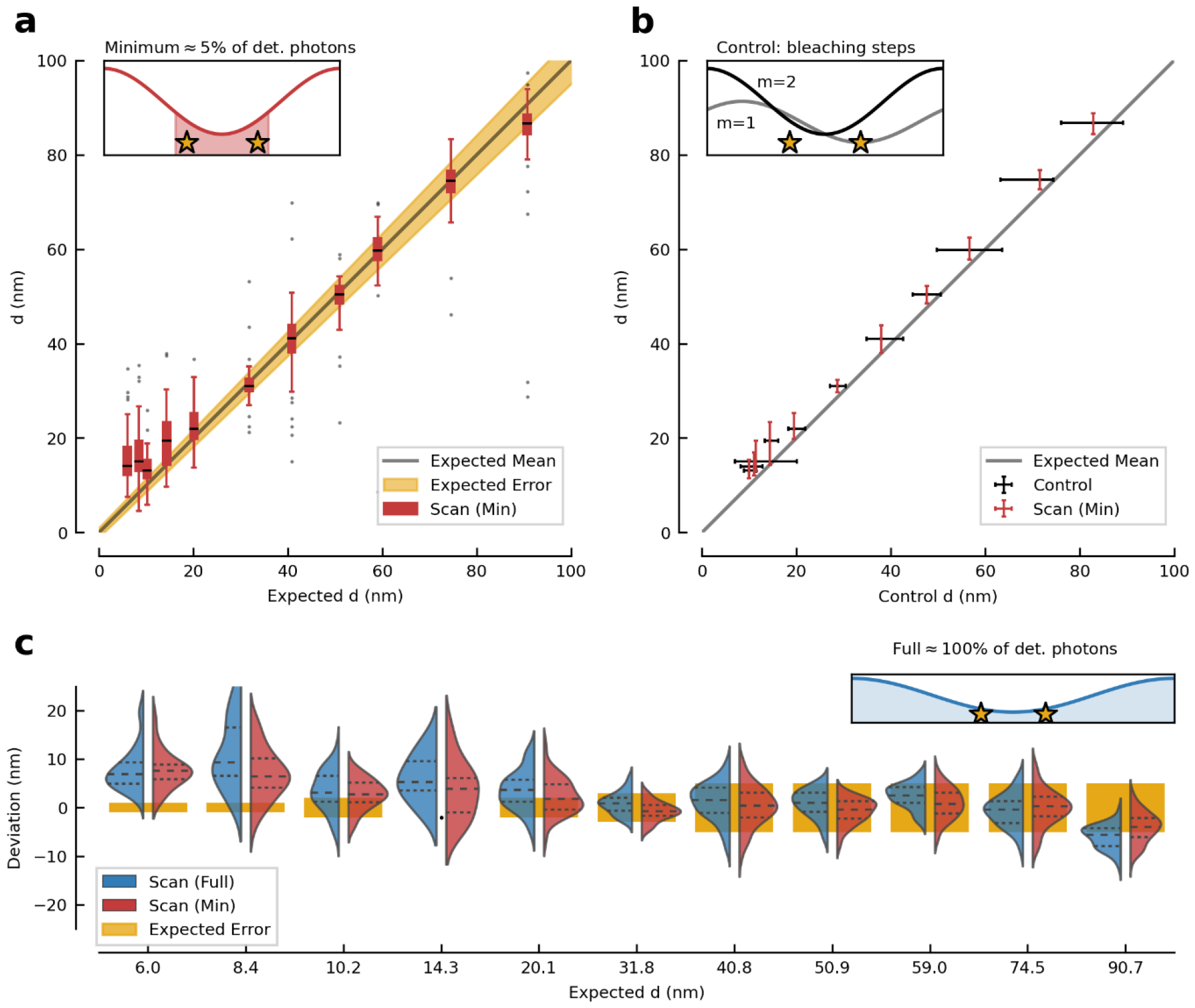
Distance measurement via line-scans and control measurement via bleaching steps. **a**: Boxplot of the distance estimates obtained from line-scan experiments with respect to expected distances up to 90 nm. Here, we determine the distance of two fluorophores by selecting photons near the minimum of a line-scan (Min, see inset). The box extends from the lower to upper quartile values of the data. Error bars mark the interval from the median (black line within box) to the last point within a 1.5 interquartile range. Outliers are shown as black dots. The experiment agrees with the expected *d* down to 10 nm. For *d*<10 nm, the estimation saturates and fails to distinguish between shorter and longer nanorulers; this can be attributed to experimental imperfections, such as a non-zero minimum intensity of the probing pattern. **b:** Correlation between obtained distance estimates and our control by centre-of-mass shift due to bleaching of one molecule. While the methods agree well on the separation estimation, our method relying on the minimum is more precise than the control and uses only a fraction of the available photons (5 %) of a full line-scan. The inset depicts how the centre-of-mass shift allows determining *d* via a bleaching step. **c**: Half-violin plot of the deviation of obtained distance estimates from the expected distance. We compare distance estimates obtained by utilizing all photons available from a full line-scan (Full, red) with estimation utilizing only photons near the emission minimum (Min, blue). Considering more photons does not improve the co-localization accuracy or precision, since photons from the maxima do not add information. On the contrary, longer measurement sequences and exposure to higher illumination intensities increase blinking of the molecules and deteriorate the measurement, for example by saturation of the emission.

