## Supplements for "Diffraction minima resolve point scatterers at tiny fractions (1/80) of the wavelength"

**Thomas Arne Hensel**[1], **Jan Otto Wirth**[2], **and Stefan W. Hell**[*1,2]

[1]Department of NanoBiophotonics, Max Planck Institute for Multidisciplinary Sciences, Am Fassberg 11, Göttingen, DE
[2]Department of Optical Nanoscopy, Max Planck Institute for Medical Research, Jahnstraße 29, Heidelberg, DE

***Correspondence:***


January 18, 2024

### 1 Calculations

In the following, we provide some more details on the derivation of the precision estimates for the multi-emitter co-localization problem in form of the Cramer Rao Bound. The setting we are interested in is depicted in Figure 1. We employ a standing wave to probe to fluorophores that are spaced far below the diffraction limit. The emission of the molecules will resemble the illumination pattern when we move the pattern with respect to the molecules coordinates.

#### 1.1 Harmonic Intensity Model

In general, the spatial profile of the emitted light is the convolution of illumination, source density and detection PSF (Gaussian). However, in our specific case, we modulate only the phase of the standing wave. This yields a perfect fringe pattern as shown in Figure 1, (right panel). Here, we fix a spatial coordinate and vary the phase of the standing wave within the Gaussian envelope, resulting in a perfect harmonic response of the system. Starting with an idealized model, we add more realistic features in the subsequent paragraphs to construct the full model eventually.

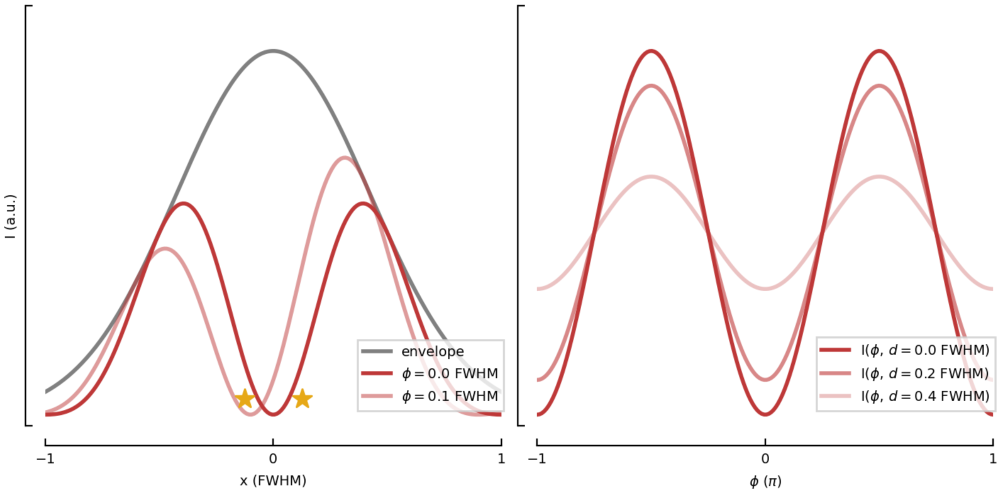

Figure 1: Emission of the molecules resembles the fringe pattern of the illumination source. Left: Spatial representation of the probing intensity profile by multiplication of a standing wave and a Gaussian envelope. By variation of the phase of the standing wave, one can adjust the position of the intensity minimum without changing the spatial position of the envelope, which results in a fringe pattern as a response of the molecules. Right: Response of the sources due to the phase modulation of the standing wave. Note that this modulation is perfectly harmonic w.r.t. the phase since the spatial position of the envelope is fixed. Depending on the position of the centre-of-mass relative to the envelope's maximum, the amplitude of the emission may vary, but it does not change its shape. The fringe contrast of this pattern depends on the distance of the sources, which allows to access information on their separation.

##### 1.1.1 Idealized Harmonic Model

Without any external disturbance, the response of the $i-th$ source to a harmonic input signal obeys

$$h(x_i, \phi) \propto \sin\left(\frac{2\pi}{\lambda} x_i + \phi\right)^2 \tag{1}$$

$$\propto \frac{1}{2}\left(1 - \cos(2(\varphi_i + \phi))\right), \tag{2}$$

where $\varphi_i = \frac{2\pi}{\lambda} x_i$ is the relative phase of the source with respect to the reference frame and $\phi$ is the relative phase of the standing wave with respect to the reference frame. The full model may be written as

$$N(\phi) = c_0 \left(h(\vec{\varphi}, \phi)\right). \tag{3}$$

$c_0$ denotes the expected photon flux for unit quantum yield of the sources. In the following, we absorb $c_0$ into the source's brightness $\gamma$ of $h(x_i, \phi)$ to consider the incident photon flux, scattering cross section of the molecules etc with a single parameter.

The combined emission $h$ as a superposition of harmonic functions with the same frequency may be rewritten as (the sum index runs from 0 to M-1 if not indicated otherwise):

$$
\begin{aligned}
h(\vec{\varphi}, \phi) =& \sum_i \gamma_i h_i(\varphi_i, \phi) && (4)\\
=& \frac{1}{2} \sum_i \gamma_i \left(1 - \cos(2(\varphi_i + \phi))\right) && (5)\\
=& \frac{1}{2} \left( \sum_i \gamma_i - \sum_i \gamma_i \cos(2(\varphi_i + \phi)) \right) && (6)\\
=& \frac{M\bar{\gamma}}{2} - \frac{1}{2} \sum_i \gamma_i \cos(\delta_i + \Phi) && (7)\\
=& \frac{M\bar{\gamma}}{2} - \frac{\Gamma}{2} \cos(\delta + \Phi) && (8)\\
=& a_0 - a_1 \cos(\delta + \Phi), && (9)
\end{aligned}
$$

where $\bar{\gamma}$ denotes the average of the sources' brightness, $\Gamma, \delta$ result as cumulative amplitude and phase from the superposition of cosines, according to the *harmonic addition theorem*:

$$
\begin{aligned}
\Gamma^2 =& \sum_i \gamma_i^2 + 2 \sum_i \sum_{j>i} \gamma_i \gamma_j \cos(\delta_i - \delta_j) && (10)\\
\tan(\delta) =& \frac{\left(\sum_i \gamma_i \cos(\delta_i)\right)}{\left(\sum_j \gamma_j \cos(\delta_j)\right)} && (11)\\
\delta_i =& 2\varphi_i = \frac{4\pi x_i}{\lambda} && (12)\\
\Phi =& 2\phi && (13)
\end{aligned}
$$

Together, $\Gamma$ and $\delta$ encode the information on the sources' positions. The amplitude $\Gamma$ of the cumulative signal of the sources (incoherent superposition of individual signal) is less or equal than the sum of the individual signals. This attenuation is paramount to co-localize the two sources, as will become clear in a moment. We can express the full model via the respective contributions to the emission:

$$
\begin{aligned}
N(\Phi) =& c_0 \left( \frac{M\bar{\gamma}}{2} - \frac{\Gamma}{2} \cos(\delta + \Phi) \right) && (14)\\
=& c_0 \left( \frac{N_{\mathrm{off}}}{c_0} - \frac{N_{\mathrm{mod}}}{c_0} \right). && (15)
\end{aligned}
$$

$N_{\mathrm{off}}$ denoting the constant offset depending on number and brightness of the sources and $N_{\mathrm{mod}}$ the (attenuated) modulation of the signal. In the following, we restrict our investigation to $M = 2$, since this is the simplest case that allows to demonstrate all the relevant physics.

###### 1.1.2 Finite Background

In order to account for a homogeneous background, e.g. due to scattering of photons from the illumination source, residual ambient light, we include a background rate $\beta$ into the model:

$$
\begin{aligned}
N(\Phi) =& c_0 \left( \beta + \frac{M\bar{\gamma}}{2} - \frac{\Gamma}{2} \cos(\delta + \Phi) \right) && (16)\\
=& c_0 \left( \frac{N_{\mathrm{bgr}}}{c_0} + \frac{N_{\mathrm{off}}}{c_0} - \frac{N_{\mathrm{mod}}}{c_0} \right). && (17)
\end{aligned}
$$

$N_{\mathrm{bgr}}$ denoting the background counts and $N_{\mathrm{off}}$, $N_{\mathrm{mod}}$ as before.

###### 1.1.3 Imperfect Intensity Minimum

Until now we have assumed a perfect standing wave as a response of the sources. In an experiment, this is usually not the case and one has to deal with imperfect fringe contrast, i.e. finite intensity minima. The noise in the minimum limits the intensity modulation one can observe, hence it puts a lower bound on the distances that can be resolved. We calculate the intensity profile of two interfering imbalanced beams with a ratio $\alpha = |E_1|/|E_0| \leq 1$ of their fields. In case of perfectly balanced interfering waves, $\alpha = 1$. For a coherent superposition of two incoming fields with phase difference $\varphi_1 - \varphi_2$, we get for the intensity in the sample plane:

$$
\begin{aligned}
I(\vec{x},\vec{\varphi}) =& |\sum_{j=0}^{1} E_j e^{i\left(\vec{k}_j\vec{x}+\varphi_j\right)}|^2 && (18)\\
=& |E_0|^2 + |E_1|^2 + E_0E_1^* e^{i\left(\vec{k}_0-\vec{k}_1\right)\vec{x}} e^{i(\varphi_0-\varphi_1)} + E_1E_0^* e^{i\left(\vec{k}_1-\vec{k}_0\right)\vec{x}} e^{i(\varphi_1-\varphi_0)} && (19)\\
=& |E_0|^2 + |E_1|^2 + |E_0||E_1| \left(e^{i\left(\Delta\vec{k}\cdot\vec{x}+\Delta\varphi\right)} + e^{-i\left(\Delta\vec{k}\cdot\vec{x}+\Delta\varphi\right)}\right) && (20)\\
=& |E_0|^2 \left(1+\alpha^2+2\alpha\cos(\Delta\vec{k}\cdot\vec{x}+\Delta\varphi)\right) && (21)
\end{aligned}
$$

Now the response $h$ of a single source has to be modified accordingly, resulting in the cumulative signal from the sources:

$$I(\vec{\varphi},\phi) = (1+\alpha^2)M\bar{\gamma} - 2\alpha\Gamma\cos(\delta(\vec{\varphi})+\phi) \quad (22)$$

We have absorbed $|E_0|$ into $\bar{\gamma}$ and set $\phi$ as the global phase of the response, which is tunable by varying the phase difference between the interfering waves.

##### 1.1.4 Unequally Bright Sources

The amplitude $\Gamma$ depends on the individual source brightness $\gamma_i$, which may be different. For two sources, we can express $\Gamma$ in terms of their average brightness and their relative difference in brightness $r$:

$$
\begin{aligned}
\Gamma^2 =& 2\bar{\gamma}^2 \left(1 + \left(\frac{\Delta\gamma}{2\bar{\gamma}}\right)^2 + \left(1 - \left(\frac{\Delta\gamma}{2\bar{\gamma}}\right)^2\right)\cos\left(\delta_1-\delta_2\right)\right) && (23)\\
=& 2\bar{\gamma}^2 \left(1 + r^2 + \left(1-r^2\right)\cos\Delta_\delta\right). && (24)
\end{aligned}
$$

Where $\bar{\gamma} = (\gamma_1+\gamma_2)/2$ denotes the average source brightness, $\Delta\gamma = |\gamma_1-\gamma_2|$ the absolute difference in brightness and $\Delta_\delta$ is the source separation in units of phase.

##### 1.1.5 Full Model

Combining all the above, the full model reads

$$
\begin{aligned}
I(\vec{\varphi},\phi) =& \bar{\gamma}\left(\beta + (1+\alpha^2)M\right) - 2\alpha\Gamma\cos(\delta(\vec{\varphi})+\phi) && (25)\\
=& a_0 - a_1\cos(\delta(\vec{\varphi})+\phi) && (26)\\
=& (N_{bgr}+N_{off}) - N_{mod} && (27)
\end{aligned}
$$

with $\Gamma$ defined as in Equation 23 and $\alpha$ as in subsubsection 1.1.3. In order to take all possible imperfections into account, we express the harmonic response in terms of the visibility. The visibility conveniently captures all different effects and provides a single dimensionless parameter to investigate the physics of the co-localization problem.

The visibility is defined as $\nu = (I_{max}-I_{min})/(I_{max}+I_{min})$, which is the quotient of modulation amplitude and average intensity.

$$\nu = a_1/a_0. \quad (28)$$

We rewrite the signal dependent on the average cumulative signal and modulation amplitude (see Equation 25) in terms of the visibility:

$$
\begin{aligned}
I(\vec{x},\vec{\varphi}) =& a_0\left(1-\nu\cos(\delta+\Delta\varphi)\right) && (29)\\
\hat{I}(\vec{x},\vec{\varphi}) =& I/\bar{I} = 1-\nu\cos(\delta+\Delta\varphi). && (30)
\end{aligned}
$$

Since we are interested in the change in visibility due to the source separation and not background, we define

$$\tilde{\nu} = a_1/(a_0 - N_{\mathrm{bgr}}) \quad (31)$$

as the background-corrected visibility. Note that this important, otherwise the visibility depends on the brightness of the sources and does not yield an informative measure of the quality of the minimum.

#### 1.2 Estimating Separation and Orientation

##### 1.2.1 Separation

We recall that the positions of the sources are reconstructed from estimates of the parameters $a_0$ and $a_1$ in Equation 25 with the help of Equation 10. For two sources in 1D, the separation reads:

$$|x_1 - x_2| = \frac{\lambda}{4\pi} \arccos\left(\frac{\Gamma^2}{2\bar{\gamma}^2} - 1\right) \tag{32}$$

We can now express the distance estimate in terms of the initial visibility $\nu_0$ and the reduced fringe contrast $\nu_1$ due to the finite distance of two sources. Wile a perfect standing wave has unit visibility, we do not presume this is the case in our experiments. Let us denote $\nu_0$ as the initial (uncorrected) visibility and $\tilde{\nu}_0$ as the background corrected initial visibility. In order to take into account possible imperfections, we assume $\tilde{\nu}_0 \leq 1$ and calculate the ratio of the systems default or initial visibility $(a_{1,0}/(a_{0,0} - N_{\mathrm{bgr}})$, known via a calibration measurement) and the observed two-source visibility:

$$\tilde{\nu}_1/\tilde{\nu}_0 = \frac{a_{1,1}}{a_{1,0}} \frac{(a_{0,0} - N_{\mathrm{bgr}})}{(a_{0,1} - N_{\mathrm{bgr}})} \tag{33}$$

$$= \frac{\Gamma_{\Delta \geq 0}}{2\bar{\gamma}} \tag{34}$$

Here $a_{1,0}$, $a_{0,0}$ denote the coefficients $a_1$, $a_0$ for the pure fringe pattern (with one molecule, two molecules with zero distance or some other point-like test object to quantify the quality of the minimum), while $a_{1,1}$, $a_{0,1}$ denote those for non-zero source separation (in the presence of two sources). The distance can now be calculated as

$$|x_1 - x_2| = \frac{\lambda}{4\pi} \arccos\left(2\left(\frac{\tilde{\nu}_1}{\tilde{\nu}_0}\right)^2 - 1\right) \tag{35}$$

This calculation is contingent on a suitable characterization of the apparatus, in order to determine the initial visibility.

If the molecules do not have the same brightness ($\gamma_1 \neq \gamma_2$), the COM does not coincide with the geometric center of the nanoruler. Measuring the offset at the minimum of the intensity profile then yields a distance that is biased due to the difference in brightness. Since the distance and the difference in brightness enter Equation 32 as a product, one cannot discern intensity fluctuations from a physically changed distance reliably. The average distance is unbiased as long as the intensity fluctuations average out over time, which is the case for laser noise, but not necessarily for molecules subject to a plethora of photo-physical effects. Note also, that the distance estimate requires a sound pre- or post-estimation of the background to be unbiased; for a low uncertainty additionally a low background is needed.

##### 1.2.2 Orientation

Given two sources with a certain separation in two dimensions, we want to determine four parameters: centre-of-mass of the system (2 parameters), the distance of the sources and the orientation of the system. While a radial symmetric illumination pattern is sufficient to retrieve information about the distance of the molecules, it does not yield information about their orientation, as I show below. In order to retrieve information about the orientation, the Fisher Information on this parameter must not be zero and hence the intensity profile has to depend on the angle $\phi_d$, specifying the orientation of the sources with respect to a local coordinate system.

The positions of the molecules $\mathrm{M}_1$ and $\mathrm{M}_2$ with respect to an arbitrary origin are given by

$$\vec{r}_1 = d_{\mathrm{COM}} \begin{bmatrix} \cos(\phi_{\mathrm{COM}}) \\ \sin(\phi_{\mathrm{COM}}) \end{bmatrix} + d/2 \begin{bmatrix} \cos(\phi_d) \\ \sin(\phi_d) \end{bmatrix} \tag{36}$$

$$\vec{r}_2 = d_{\mathrm{COM}} \begin{bmatrix} \cos(\phi_{\mathrm{COM}}) \\ \sin(\phi_{\mathrm{COM}}) \end{bmatrix} + d/2 \begin{bmatrix} \cos(\phi_d + \pi) \\ \sin(\phi_d + \pi) \end{bmatrix} \tag{37}$$

Here $d_{\mathrm{COM}}$ denotes the distance from the origin to the centre-of-mass of the molecules, $\phi_{\mathrm{COM}}$ denotes the angle with respect to an arbitrarily chosen orientation of the coordinate system, defined by the illumination pattern. Similarly, $d$ and $\phi_d$ denote the distance and orientation of the molecules in their local coordinate system, attached to their centre-of-mass. The orientation of which is the same as that of the global coordinate system (see Figure 2).

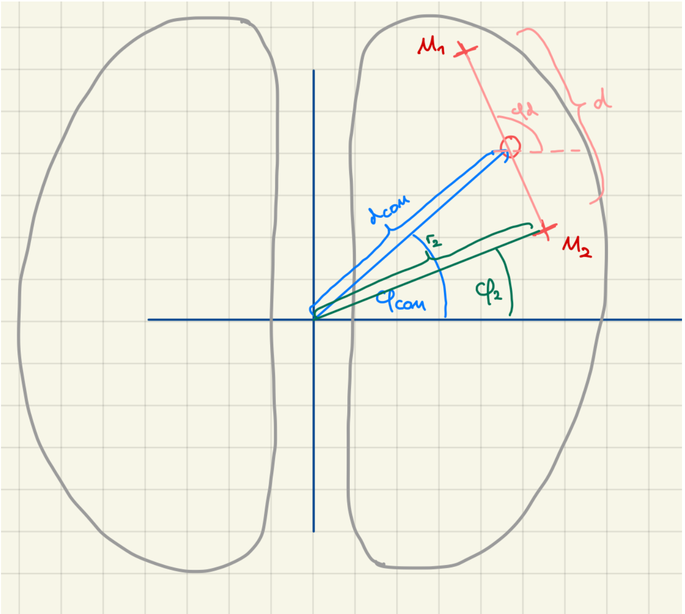

Figure 2: Schematic of the orientation problem and nomenclature of the used angles and distances. $M_1$ and $M_2$ denote the two molecules.

The distance $r_1$ to the origin is given by

$$r_1(d_{\text{COM}}, d, \phi_{\text{COM}}, \phi_d) = \sqrt{d_{\text{COM}}^2 + (d/2)^2 + d_{\text{COM}} d \cos(\phi_{\text{COM}} - \phi_d)/2}. \tag{38}$$

Hence $r_2 = r_1(d_{\text{COM}}, d, \phi_{\text{COM}}, \phi_d + \pi)$. We now compare an illumination beam profile that is radially symmetric with one breaking the symmetry. For example, one could think of patterns that imitate a $\text{TEM}_{01*}$-mode shape (doughnut) and $\text{TEM}_{01}$ mode with a line-shaped minimum. Both exhibit approximately quadratic intensity profiles near the minimum, one without a dependence on the orientation, the other with an angular dependence. Superposing the two fluorescence signals incoherently yields the following intensity patterns at the detector:

$$I_{01*} \propto r_1^2 + r_2^2 \tag{39}$$

$$= 2\left(d_{\text{COM}}^2 + \left(\frac{d}{2}\right)^2\right) \tag{40}$$

$$I_{01} \propto r_1^2 \cos(\phi_1) + r_2^2 \cos(\phi_2) \tag{41}$$

$$= \left(d_{\text{COM}}^2 + \left(\frac{d}{2}\right)^2\right)\left(1 + \cos(\tilde{\alpha})\cos(\tilde{\beta})\right) - d_{\text{COM}}\frac{d}{2}\cos(\phi_{\text{COM}} - \phi_d)\sin(\tilde{\alpha})\sin(\tilde{\beta}), \tag{42}$$

where $\tilde{\alpha} = 2d_{\text{COM}}\cos(\phi_{\text{COM}})/r_1$ and $\tilde{\beta} = d\cos(\phi_d)/r_1$. The superposition of the doughnut signals is already independent of $\phi_d$, which inhibits the retrieval of orientation-related information. Evaluating the second expression for the general situation of a pre-localized system (i.e. illumination pattern pointing at the effective center-of-mass of the sources) allows to retrieve the desired information on the orientation, even for unequally bright sources. For this scenario, we have $d_{\text{COM}} = 0$, $r_1 = d_1$, $r_2 = d_2$ and

$$I_{01} \propto \left(\frac{d}{2}\right)^2 \left(1 + \cos\left(\frac{d}{d_1}\cos(\phi_d - \phi_{\text{PSF}})\right)\right). \tag{43}$$

By varying the orientation of the PSF $\phi_{\text{PSF}}$, one can extract the angle $\phi_d$. With two perpendicular orientations one can efficiently extract the rulers orientation, essentially by projecting the distance in the direction of the exposure (e.g. x and y). This leaves only slight ambiguities, since permutations of the molecules cannot be distinguished. A third exposure would be necessary to couple both axes and make the result unique. A radially symmetric illumination profile does not yield any information about the orientation of the two sources.

The latter can also be obtained as a simple trigonometric fact (see Figure 3). Any triangle that has a median length of $m_a$ with respect to the side $a$ and where $a$ lies on a circle (blue), has a fixed

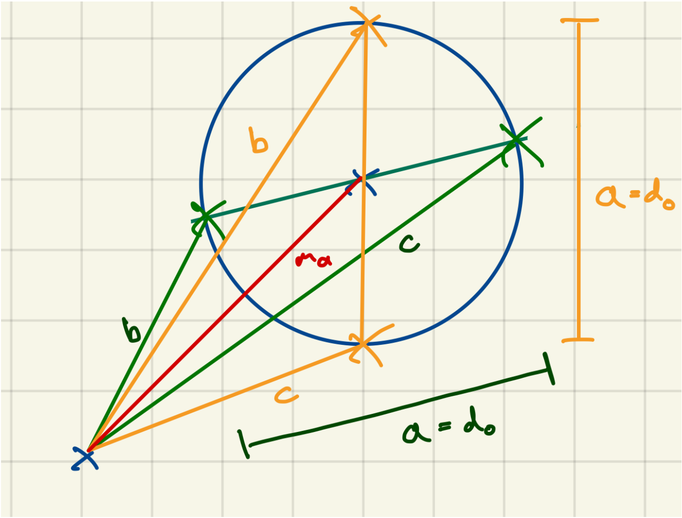

Figure 3: The sum of the square of the sides of any triangle with the same length a and median $m_a$ is constant.

sum of the quadratures of the sides b and c, which corresponds to the incoherent superposition of the individual signals. Therefore, the resulting intensity is independent of the orientation:

$$m_a = \frac{\sqrt{2(b^2+c^2)-a^2}}{2} \tag{44}$$

$$\implies \frac{2m_a^2+a^2}{2} = b^2+c^2 \tag{45}$$

$$\implies \frac{2d_{\mathrm{COM}}^2+d_0^2}{2} = b^2+c^2 \tag{46}$$

$$= const. \tag{47}$$

#### 1.3 Fisher Information and Cramer Rao Bound

This section introduces the information theoretical framework employed for a maximum likelihood estimation of source separation. In coordinate-targeted methods one probes the sample with an illumination source exhibiting some spatial structure. The sample interacts with the probing beam and emits fluorescent photons. The probing positions and respective photon counts are recorded. If the illumination pattern and the characteristics of the stochastic behaviour of the emission are known, one can infer the most likely parameters of the underlying emitter density. We refer to the illumination pattern as the *model* $\lambda$ and the set of *parameters* $\{p\}$ parametrizes the emitter density (e.g. the location and brightness of a point source etc).

##### 1.3.1 General Framework

Now let $\{\theta\}$ be a set of parameters of a function $\lambda(\{\theta\})$. Let $P(\lambda)$ be an inhomogeneous poisson distribution with mean $\lambda(\{\theta\})$, such that $P(\lambda, n) = \lambda^n e^{-\lambda}/n!$, i.e. the probability for outcome $n$ with an underlying mean $\lambda$ is poisson distributed and given by $P$. In our case $n$ is a photon number and we wish to determine the parameters $\{\theta\}$ via the observable instances of our model $\lambda_j(\{\theta\})$, where the index $j$ denotes the respective realization of the inhomogeneous Poisson process. The probability to measure a certain ordered set of photon counts is given by:

$$P\left(\{n\}|\lambda(\{p\})\right) = \prod_j \frac{\lambda_j^{n_j} e^{-\lambda_j}}{n_j!} \tag{48}$$

$$= e^{-\sum_j \lambda_j} \prod_j \frac{\lambda_j^{n_j}}{n_j!} \tag{49}$$

Here $j$ denotes a certain experimental configuration, that is the current state of the underlying model $\lambda$ as instance of the inhomogeneous Poisson process. $\lambda_j$ denotes the expected model mean in photon counts, while $n_j$ is the respective (measured) count in $\{n\}$. $P$ is called the likelihood function. The elements of the Fisher Information matrix wrt parameter $\theta$ are given by

$$[I(\{\theta\})]_{ij} = E[(\partial_{\theta_i} \log P(\lambda(\theta))(\partial_{\theta_j} \log P(\lambda(\{\theta\}))] \tag{50}$$
$$= -E[\partial_{\theta_i}\partial_{\theta_j}\mathcal{L}(\lambda(\{\theta\})] \tag{51}$$

$\mathcal{L}(\lambda(\{\theta\})$ denotes the so-called log-likelihood. For a poissonian process, it is given by:

$$\mathcal{L}(\{n\}|\lambda(\{\theta\}) = \log P \tag{52}$$
$$= -\sum_j \lambda_j + \sum_j \log \frac{\lambda_j^{n_j}}{n_j!} \tag{53}$$
$$= -\sum_j \lambda_j + \sum_j \log \lambda_j^{n_j} - \log n_j! \tag{54}$$
$$= \sum_j -\lambda_j + n_j \log \lambda_j - \log n_j! \tag{55}$$

Its first and second derivatives are calculated as

$$\partial_{\lambda_k}\mathcal{L}(\{n\}) = \sum_j n_j \frac{1}{\lambda_j}\delta_{jk} - \delta_{jk} \tag{56}$$
$$= \delta_{jk}\left(\sum_j \frac{n_j}{\lambda_j} - 1\right) \tag{57}$$
$$= \frac{n_k}{\lambda_k} - 1 \tag{58}$$
$$\partial_{\lambda_m}\partial_{\lambda_k}\mathcal{L}(\{n\} = \frac{n_k}{\lambda_k^2}\delta_{km} \tag{59}$$

Therefore:

$$[I(\{\bar{n}\})]_{mk} = -E[\partial_{\lambda_m}\partial_{\lambda_k}\mathcal{L}(\{\bar{n}\})] \tag{60}$$
$$= \frac{1}{\lambda_m}\delta_{km}. \tag{61}$$

The Fisher Information matrix is diagonal in the basis of the average model values (means of the poisson process) with shape $K \times K$, $K$ denoting the number of measurements taken. The Cramer Rao Bound yields a lower bound for any asymptotically unbiased and efficient estimator. In the case of multiple (spatial) parameters, we define the average Cramer Rao Bound as

$$\sigma_{\mathrm{CRB}} = \sqrt{(\mathrm{Tr}(I^{-1})/\#\{\theta\}} \tag{62}$$

We apply this formalism to two problems. On the one hand, we are interested in the Fisher Information matrix $I$, since the eigenvalues of $I^{-1}$ yield the variances of the model values $\lambda_j$. Eventually, we want to know $\sigma(\{\theta\})$ instead of $\sigma(\{\lambda\})$. We therefore need to transform $I$ and find a simple expression for its inverse. On the other hand, the (log)likelihood function allows to find the most probable model parameters by means of optimization. Minimization of the negative (log)likelihood yields a so-called *Maximum Likelihood Estimate.* We later employ this idea to find the most likely model parameters, i.e. source positions.

##### 1.3.2 Covariances In Parameter Coordinates

In order to find the covariances of the model parameters, we apply the following routine:

1. Calculate $I$ in the $\{\lambda_j\}$ (model mean) basis.
2. Perform a coordinate transformation into the $\{\theta_j\}$ (model parameter) basis.
3. Perform a basis transformation to the $\{\tilde{\theta}_j\}$ basis, e.g. in order to find coordinates that diagonalize $I$.[1]
4. Invert $I$ in this transformed state to retrieve the variances as Eigenvalues of $I^{-1}$.

[1]This could be centre-of-mass and separation instead of cartesian coordinates of the molecules positions.

Transforming $I$ into suitable parameter coordinates is done by applying a the respective Jacobian matrix and a basis transformation:

$$
\begin{aligned}
I(\{\theta\}) =& J^T I(\{\lambda\}) J && (63)\\
I(\{\tilde{\theta}\}) =& T_{B'}^{B} J^T I(\{\lambda\}) J T_{B}^{B'} && (64)\\
I^{-1}(\{\tilde{\theta}\}) =& {T_{B}^{B'}}^{-1} (J^T I(\{\lambda\}) J)^{-1} {T_{B'}^{B}}^{-1} && (65)\\
=& T_{B'}^{B} (J^T I(\{\lambda\}) J)^{-1} T_{B}^{B'} && (66)
\end{aligned}
$$

The Jacobi matrix is defined as

$$
[J]_{ij} = \partial_j \lambda_i. \tag{67}
$$

The elements of the transformed Fisher Information matrix are given by:

$$
\begin{aligned}
[J^T I J]_{ij} =& \sum_{m=1}^{l} \left( \sum_{k=1}^{n} \frac{\partial \lambda_k}{\partial \theta_i} \lambda_k^{-1} \right) \frac{\partial \lambda_m}{\partial \theta_j} && (68)\\
=& \left( \sum_{k=1}^{n} (\partial_i \lambda_k) \lambda_k^{-1} \right) \left( \sum_{m=1}^{l} \partial_j \lambda_m \right) && (69)\\
=& \sum_{k=1}^{n} (\partial_i \lambda_k) \left( \sum_{m=1}^{l} \partial_j \lambda_m \right) \lambda_k^{-1} && (70)\\
=& \sum_{k,m}^{n,l} (\partial_i \lambda_k)(\partial_j \lambda_m) \lambda_k^{-1} && (71)
\end{aligned}
$$

The analytical inversion of this matrix and subsequent determination of Eigenvalues is generally quite involved. However, since the above is a Hermitian positive-definite matrix, one can apply a Cholesky decomposition to efficiently invert the matrix and then determine its eigenvalues. For a simple one dimensional case, we can find an analytical expression for the Cramer Rao Bound, as shown below.

##### 1.3.3 CRB comparison for maximum and minimum

We compare the CRB of the distance estimate for two scenarios: measuring with an intensity maximum and an intensity minimum. To that end, we calculate the Fisher Information for a three-point measurement ($\Phi \in \{\pm L/2, 0\}$) for the model given in Equation 29:

$$
\begin{aligned}
I(\Phi) =& \bar{\gamma}(1 - \nu(\Delta x) \cos(\Phi + \Phi_0)) && (72)\\
=& \bar{\gamma}(1 + (-1)^k \nu(\Delta x) \cos(\Phi)) && (73)
\end{aligned}
$$

with $\Phi_0 = \pi$ for the maximum (k is even) and $\Phi_0 = 0$ for the minimum (k is odd). With respect to the comments above, we begin in a basis of cartesian coordinates and transition to a basis of separation and centre-of-mass, which diagonalizes the Fisher Information Matrix. For a single parameter of interest, the Fisher Information Matrix has a single element given by:

$$
F = \sum_{i=1}^{2} \frac{1}{I(\Phi_i)} \left( \frac{\partial I(\Phi_i)}{\partial \Delta x} \right)^2. \tag{74}
$$

To express the Fisher Information in terms of photons N instead of the brightness $\bar{\gamma}$, we substitute the brightness via:

$$
\bar{\gamma} = N / \sum_i I(\Phi_i) \tag{75}
$$

Since $I(+\Phi) = I(-\Phi)$ and $(\partial_{\Delta x} I(+\Phi))^2 = (\partial_{\Delta x} I(-\Phi))^2$, we can simplify our calculation and get

$$
\begin{aligned}
\sigma_{\mathrm{CRB}}(\Delta x) =& 2\sqrt{2} \left( \frac{(-1)^{-2k} (\cos(\Delta \mathrm{x}) + 1) \csc^2(\Delta \mathrm{x})}{\nu_0^2 N} \times \right.\\
& \left. \times \frac{\left( \frac{(-1)^k \nu_0 \sqrt{\cos(\Delta \mathrm{x}) + 1}}{\sqrt{2}} + 2 \left( \frac{(-1)^k \nu_0 \sqrt{\cos(\Delta \mathrm{x}) + 1} \cos\left(\frac{L}{2}\right)}{\sqrt{2}} + 1 \right) + 1 \right)}{\left( \frac{1}{(-1)^k \nu_0 \sqrt{\cos^2\left(\frac{\Delta \mathrm{x}}{2}\right)} + 1} + \frac{2(\cos(L) + 1)}{\sqrt{2}(-1)^k \nu_0 \sqrt{\cos(\Delta \mathrm{x}) + 1} \cos\left(\frac{L}{2}\right) + 2} \right)} \right)^{1/2} && (76)
\end{aligned}
$$

This is the full analytical expression of the Cramer Rao bound. We develop this expression in a Taylor series for small $\Delta x$ and in the limit of infinitesimal probing range $L$:

$$\lim_{L\to 0} \sigma_{\mathrm{CRB}}(\Delta x) = \frac{24(-1)^k\nu_0 + 24}{6\Delta x \nu_0 \sqrt{N}\operatorname{sgn}\left((-1)^k\nu_0 + 1\right)} + \frac{\Delta x\left(1 - 2(-1)^k\nu_0\right)}{6\nu_0\sqrt{N}\operatorname{sgn}\left((-1)^k\nu_0 + 1\right)} + O\left(\Delta x^3\right) \tag{77}$$

In the limit of unit initial visibility (no imperfect fringes, no background), this yields:

$$\lim_{\nu_0\to 1} \sigma_{\mathrm{CRB,min}} = \frac{\Delta x}{2\sqrt{N}} \tag{78}$$

$$\lim_{\nu_0\to 1} \sigma_{\mathrm{CRB,max}} = \frac{8}{\Delta x\sqrt{N}} - \frac{\Delta x}{6\sqrt{N}} \tag{79}$$

We recover the well-known divergent behaviour of the CRB for the separation estimation with a maximum and a surprising linear scaling for estimation with the minimum. The latter yields a constant relative error, which allows in principle diffraction-unlimited separation estimation.

#### 1.4 Estimators

##### 1.4.1 Polynomial Estimator

A Taylor expansion of the harmonic model in Equation 25 near an extremum yields a polynomial ansatz to retrieve the sought-for parameters $\Gamma$ and the phase $\delta$. We can either expand the intensity profile around the maximum or the minimum. The PSF profile can be approximated via

$$N(\Phi) = a_0 \pm a_1 \left(-1 + \frac{(\Phi - \delta)^2}{2} - \frac{(\Phi - \delta)^4}{24} + R_6\right)$$

up to fourth order, with a 6-th order error given by $R_6$. Depending on whether we are interested in the maximum or minimum, we choose the appropriate sign (+ for the minimum, - for the maximum) $a_0$ and $a_1$ are defined as in Equation 25. This allows us to fit the curvature $a_1$ and the offset $a_0 - a_1$ to the data in a least-squares sense. For a known background, the distance of the molecules can be estimated from $a_0$ and $a_1$ under help of Equation 32.

Note that the offset and the curvature are not independent. Both depend on $a_1$ and hence on the distance as well as the average brightness. However, since $a_0$ only depends on $\bar{\gamma}$, we can eliminate the dependence of $a_1$ on $\bar{\gamma}$ by normalizing on $a_0$ (approximately for equally bright molecules) and calculate $\Delta x$ independent of the brightness.

##### 1.4.2 Harmonic Maximum Likelihood Estimator

Here, we consider the full harmonic model as in Equation 25 and perform a maximum likelihood fit of the model parameters to the data. This directly yields $a_0$ and $a_1$ and allows for a most accurate consideration of the fringe profile over the full (or partial) line scan. The maximum likelihood estimation formally follows the approach described in subsection 1.3.

##### 1.4.3 Fourier Analysis

In this approach, we extract the amplitude $a_1$ and eventually $\Gamma$ as well as the phase shift $\delta$ from $N_{\mathrm{mod}}$ in Equation 25 via a Fourier transformation. Lets decompose $N_{\mathrm{mod}}$ into a noise-free modulation part (signal) and a noise term. The Fourier transformation of $N_{\mathrm{mod}}$ is then given by:

$$\mathscr{F}(N_{\mathrm{mod}}) = \mathscr{F}(N_{\mathrm{sig}}) + \mathscr{F}(N_{\mathrm{noise}}) \tag{80}$$

$$= a_1\left(e^{i\delta}\mathscr{F}(e^{i\phi}) + e^{-i\delta}\mathscr{F}(e^{-i\phi})\right) + \mathscr{F}(N_{\mathrm{noise}}) \tag{81}$$

$$= a_1\left(e^{i\delta}\sqrt{2\pi}\delta(\omega-\omega_0) + e^{-i\delta}\sqrt{2\pi}\delta(\omega+\omega_0)\right) + \mathscr{F}(N_{\mathrm{noise}}) \tag{82}$$

$$= a_1\sqrt{2\pi}\left(e^{i\delta}\delta(\omega-\omega_0) + e^{-i\delta}\delta(\omega+\omega_0)\right) + \mathscr{F}(N_{\mathrm{noise}}) \tag{83}$$

The back transform of this expression restricted to $\omega = \pm\omega_0$ yields $\Gamma$ and $\delta$, while effectively filtering out the Poissonian noise, since the amplitude modulation of the line scan occurs at much lower frequency than the poisson noise of the emission or detector noise. Similarly, the constant offsets in Equation 16, $N_{\mathrm{bgr}} + N_{\mathrm{off}}$ yield a constant contribution at $\omega = 0$. By effectively creating a narrow band-pass filter, we eliminate the high-frequency poisson noise and zero-frequency constant offsets:

$$\mathscr{F}^{-1}(\mathscr{F}(N)\big|_{\omega=\pm\omega_0}) = \mathscr{F}^{-1}\left((\mathscr{F}(N_{\text{bgr}} + N_{\text{off}})) - \mathscr{F}(N_{\text{sig}}) - \mathscr{F}(N_{\text{noise}}))\,\big|_{\omega=\pm\omega_0}\right) \tag{84}$$

$$= - N_{\text{sig}} \tag{85}$$

$$= - a_1 \cos(\delta + \Phi), \tag{86}$$

Which is exactly the term of interest. Since

$$\mathscr{F}(N_{\text{bgr}} + N_{\text{off}}) = \sqrt{2\pi}\left(\bar{N}_{\text{bgr}} + \bar{N}_{\text{off}}\right)\delta(\omega - 0), \tag{87}$$

We can filter out background and offset as constants appearing at zero frequency. The poissonian noise of these contributions is filtered away with the same argument as the noise of the modulation. In the above, $\delta(\omega - 0)$ denotes a delta distribution.

##### 1.4.4 Correlative analysis

The signal in Equation 25 is represented by a harmonic function with a fixed frequency, amplitude, and phase. We consider the noisy signal as an instance of a random variable and a collection thereof as a vector of random variables. We know the wavelength of the fringe pattern/standing wave and now compute the correlation between the data $\vec{y}$ and a harmonic waveform template $\hat{y}$. $\vec{y}$ has a fixed but unknown relative phase shift $\phi_0$ w.r.t. the template. We extract amplitude and phase of the data as follows:

$$\vec{y}(\phi) = a \exp(i\vec{\phi}) \tag{88}$$

$$\hat{y}(\phi) = \exp(i(\vec{\phi} + \phi_0)) \tag{89}$$

$$z \equiv \langle \vec{y}, \hat{y} \rangle / \mid \hat{y} \mid \tag{90}$$

$$\mid z \mid = a \tag{91}$$

$$\Im(\log(z)) = \phi_0 \tag{92}$$

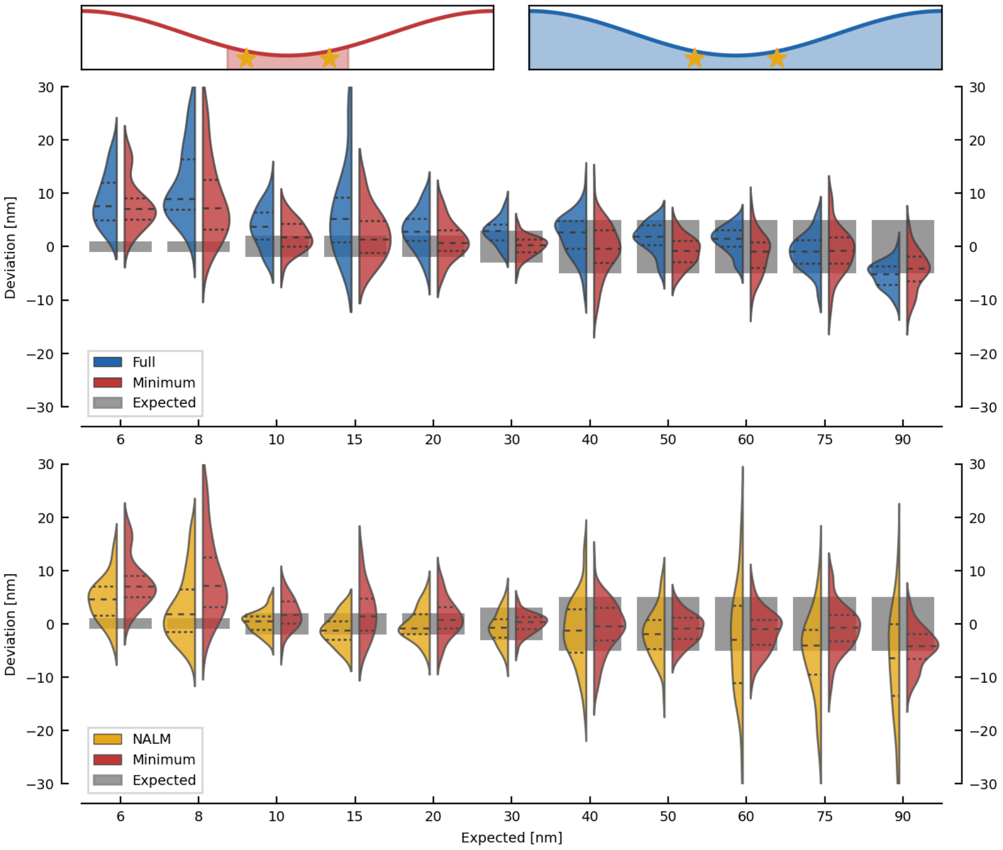

Figure 4: Resolving distances down to 10 nm with 3% of the total number of photons from a full line scan. Top: Sampling with a minimum (left) and doing a full line scan (right). Middle: Deviation of the distance estimates from the expected value w.r.t. ground truth (gray patch in the background marking the uncertainties as provided by Gattaquant) when considering photons from the minimum (red violins) or from the full line scan (blue violins). Within the violins, thick dashed lines mark the median and thin dashed lines the second and third quartile. Outliers beyond the 1.5 interquartile range from the median are not shown. Photons from the minimum suffice to estimate the distance of the sources as good as when using all available photons. Bottom: Comparison of our method and distance estimation via bleaching steps as control. The deviation w.r.t. ground truth exhibits similar behaviour for both control and our method.

### 2 Line Scan Analysis

#### 2.1 Introduction

Our theoretical model implies that two sources can be co-localized more precisely when probed with a minimum instead of a maximum. We therefore set up an experiment to compare the resolution capabilities of photons collected at the maximum and at the minimum. For maximum comparability, we wish to collect photons from the same sample. Therefore, we perform a $2\pi$ line scan on two sources with a certain distance d by scanning a fringe pattern of a standing wave over the structure[2]. Our system is a so called *Nanoruler* produced by the company Gattaquant [2]. This is a rigid DNA origami structure, which carries two fluorophores with a well defined separation. Figure 1 depicts the line scan principle. Each source (molecule on the Nanoruler) will resemble the illumination fringe pattern. The key idea is that the superposition of two fringe patterns modulates the fringe contrast if they are shifted relative to each other. Hence, a change in distance of the fluorophores appears as a change in visibility of the pattern and we can compare the performance of photons from the maximum and from the minimum to detect small changes in visibility. The results of this comparison are shown in Figure 4 and indicate that photons from the minimum suffice to co-localize two sources well below the diffraction limit.

[2] For a comprehensive overview of the experimental setup, the interested reader is referred to [1]

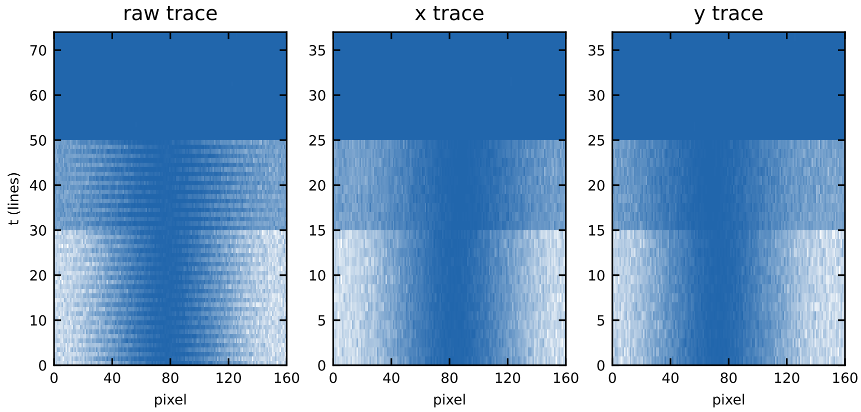

Figure 5: Raw trace with consecutive line scans. The trace is split into x- and y-axis for further processing.

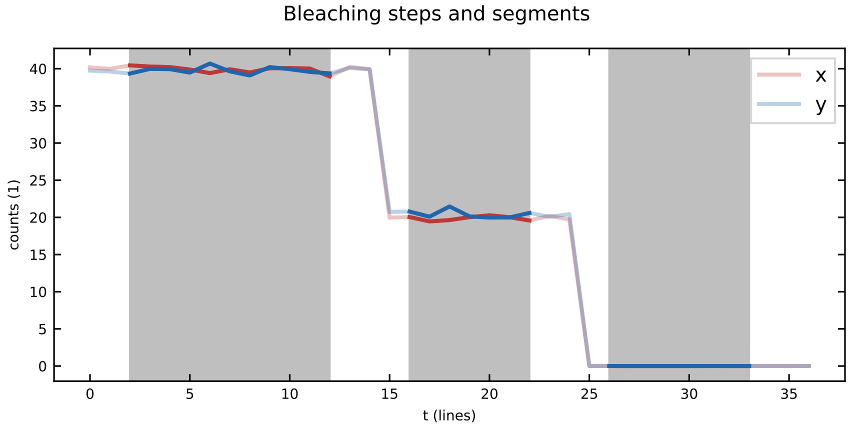

Figure 6: Trace segmentation. The trace is segmented at the bleaching steps via change point detection on the line-averaged signal. A fraction of each segment that has minimum variance within the segment is chosen for analysis.

#### 2.2 Data analysis

In the following, more details on the processing of the line scans are provided. A line scan measurement consist of a number of consecutive scans in x- and y-direction that are stacked into one image file. By varying the phase of the illumination pattern from 0 to $2\pi$ in steps of 2 nm, we obtain lines with 160 pixels. Usually 200 to 400 lines form one image.

The images are split into x- and y-scans, since the scan directions are independent of each other (see Figure 5 and [1] for details of the setup). Accordingly, we process both directions independently to obtain a distance estimate in each axis and calculate the final distance as their norm. In order to do so, we take the x- and y-scans and apply a line-wise average in each axis. This allows to detect the bleaching steps via rapid changes in brightness w.r.t. time. We use the Python package *ruptures* with a piece-wise constant model and a penalized segmentation routine (PELT) to detect a dynamic number of change points [3].

The trace is segmented at the detected change points, while we truncate each segment by a five lines before and after the bleaching step to ensure a stable state after the transition (see Figure 6). Ideally, the trace has three segments. In case of more/less molecules, bleaching steps or significant changes in brightness over the course of the measurement, there will be more segments.

Next, we filter the traces to ensure that we pick only those with two bleaching steps, i.e. three segments. This is necessary to ensure that our signal stems from two molecules. However, our method only requires the signal from the two-molecule part of the trace, as will be explained later. Additional filtering is performed on the segments to restrict the standard deviation of the average counts to less than 10 times the expected poisson noise of the line's mean. The ratio of counts between the two molecule and the one molecule segment is allowed to deviate $\pm 50\%$. While the latter two criteria are quite relaxed, the number of segments is crucial to sort out invalid measurements. This filter has a success rate of 96% on data that has been pre-selected by eye in

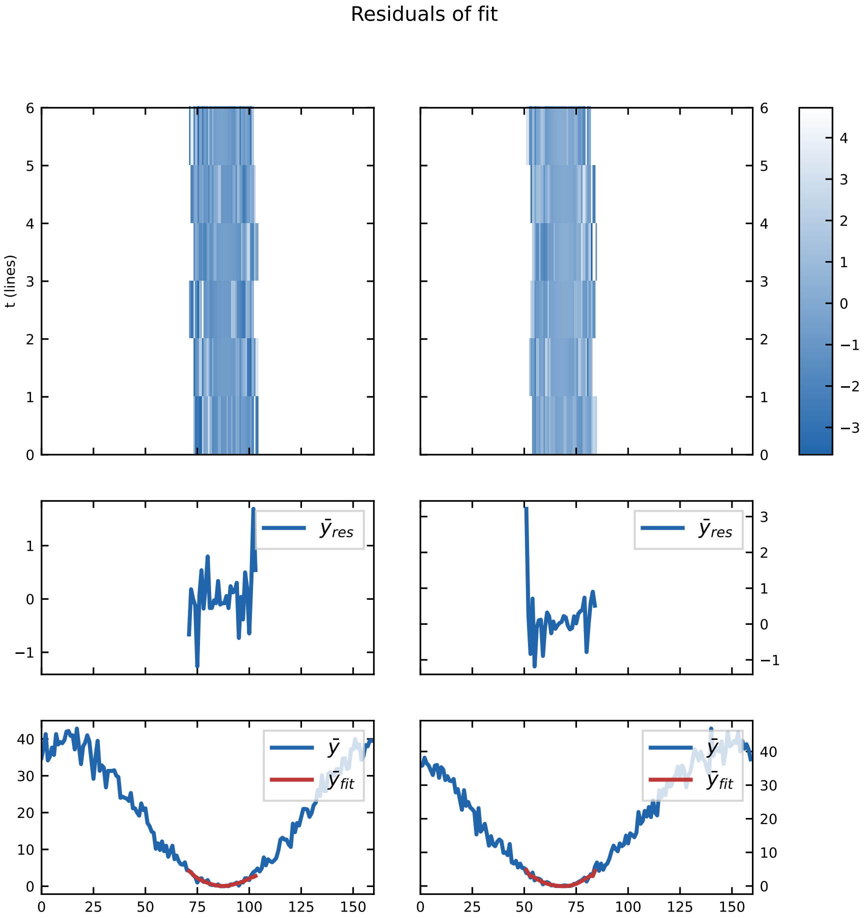

Figure 7: Local fit of a polynomial ansatz to the minimum of the harmonic intensity profile. First row: residuals of the fit wrt the individual lines that were fitted (y) and the pixel index (x). Middle row: line-averaged residuals wrt pixel index. Bottom row: Overlay of line-averaged data and fit.

order to discard obviously faulty measurements, e.g. as in the case of a cluster of molecules or a lack of bleaching steps, i.e. if only one molecule was present or the two molecules did not bleach consecutively during the measurement.

Once the filtering is complete, we transit to the evaluation of the line scans. Since the quality of the intensity minimum as well as the background may change within the sample, we decided to perform a calibration for each line scan by estimating the local background from the corresponding segment of the trace and the quality of the intensity minimum (initial fringe contrast) from the single molecule segment. This estimate serves as calibration for the following evaluation of the two molecule segment. The two molecule segment is evaluated either globally by averaging all lines in an axis or locally (i.e. each individual line or in a bootstrapping approach) with a range of methods.

In essence, we extract the parameters $a_0$ and $a_1$ from the cumulative signal (see Equation 25) There is a range of options, such as polynomial fit near the intensity minimum (see Figure 7), a maximum likelihood optimized fit of a harmonic model on the full $2\pi$ scan, a Fourier transformation accompanied with filtering for the prevalent frequencies or a correlative analysis by correlating the signal with a complex harmonic template (see subsection 1.4).

All of these methods yield an estimate of $a_0$ and $a_1$, which allows to calculate a distance estimate (projection of the distance in the respective axis) according to Equation 32. As a control method, the distance of two molecules was also obtained via the centre-of-mass shift after the first bleaching step (see subsection 2.3. The methods agree very well on the experimental data (see Figure 8), but the polynomial ansatz in the vicinity of the minimum yields the most precise and robust distance estimate with the fewest photons.

In summary, the fully automatized analysis routine proceeds in several stages:

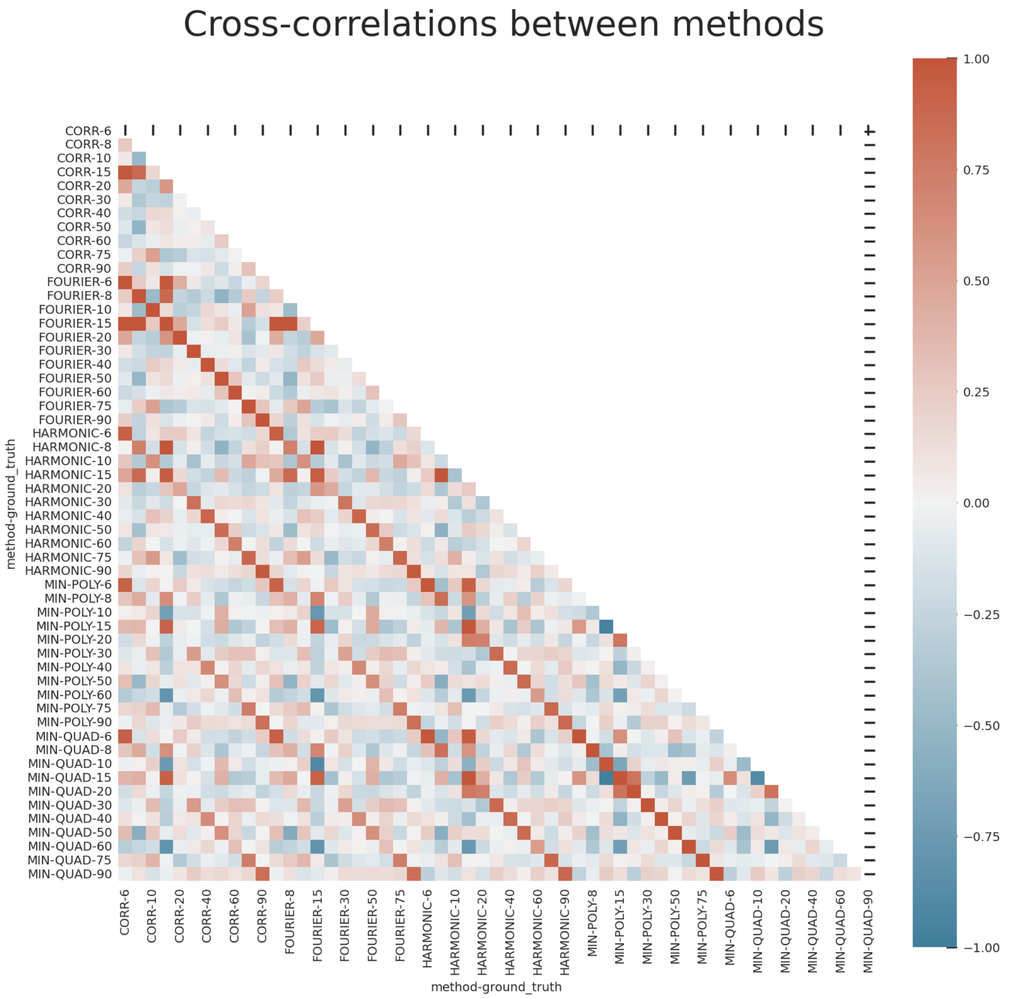

Figure 8: Correlation between estimators. The experimental data was analyzed with all estimators described in subsection 1.4. This correlation matrix quantifies the agreement of the estimators across the different sizes of Nanorulers. The agreement across the methods is apparent by the periodic diagonal signatures in the matrix, while all methods fail to distinguish sub-10 nm distances - hence the results are more random and correlation decreases for these sizes.

- Data import and splitting of trace into axes.
- Segmentation of trace wrt bleaching steps: extract two, single and zero molecule signals.
- Filtering of traces according to number of segments, variance of count rate and ratio of average counts among segments.
- Calibration: estimate for background and residual intensity in the minimum (initial visibility) on the single molecule trace.
- Global or local fit of a model to the individual or averaged lines of the respective segments.
- Application of a suitable estimator on the fit results under consideration of the calibration information.
- Joining the results of each axis to obtain the norm of the distance.

In the following, we show the distribution of obtained distance estimates in some more detail (see **??**. We plot a histogram of the obtained distances for each batch (size of Nanoruler) along with the cumulative probability density and a correlation plot of our estimate with the control. The histogram and the cumulative probability provide information about the median and the width of the distribution, while the correlation plot visualizes the (dis-)agreement with our control method. For 6 nm and 8 nm, the distances cannot reliably be estimated and the corresponding distributions are too wide and have a shifted mean in comparison to sizes larger than 10 nm. We observe that our method is not only more efficient with respect to photons needed for the estimate, but also more precise than the control, especially for larger distances.

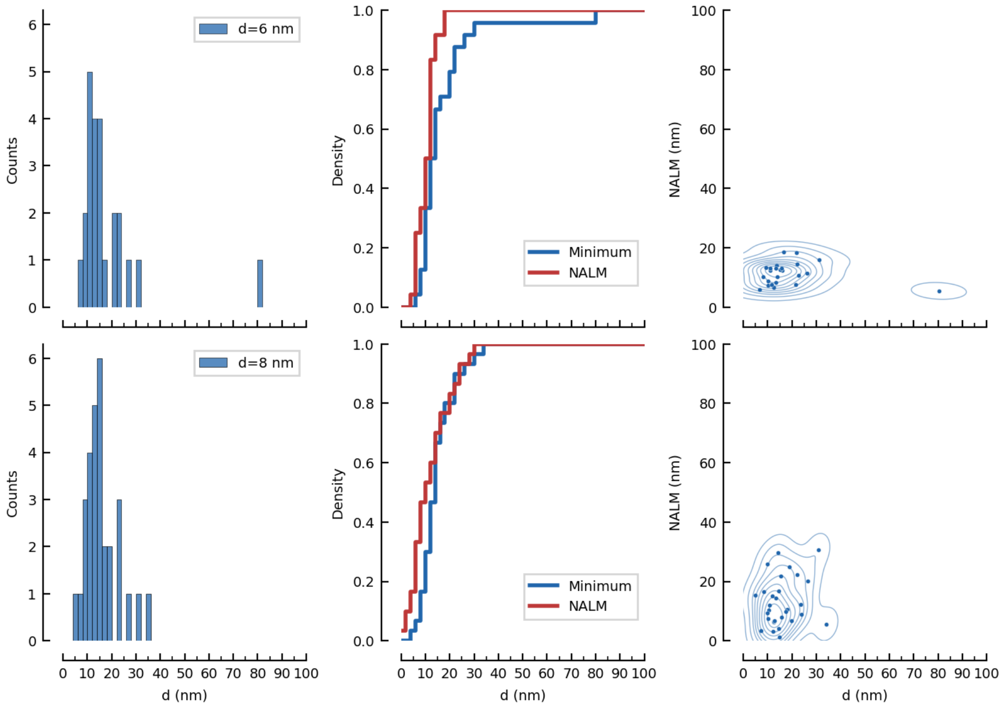

Figure 9: Statistical information on the distance estimation from line scans as described above for 6 nm and 8 nm.

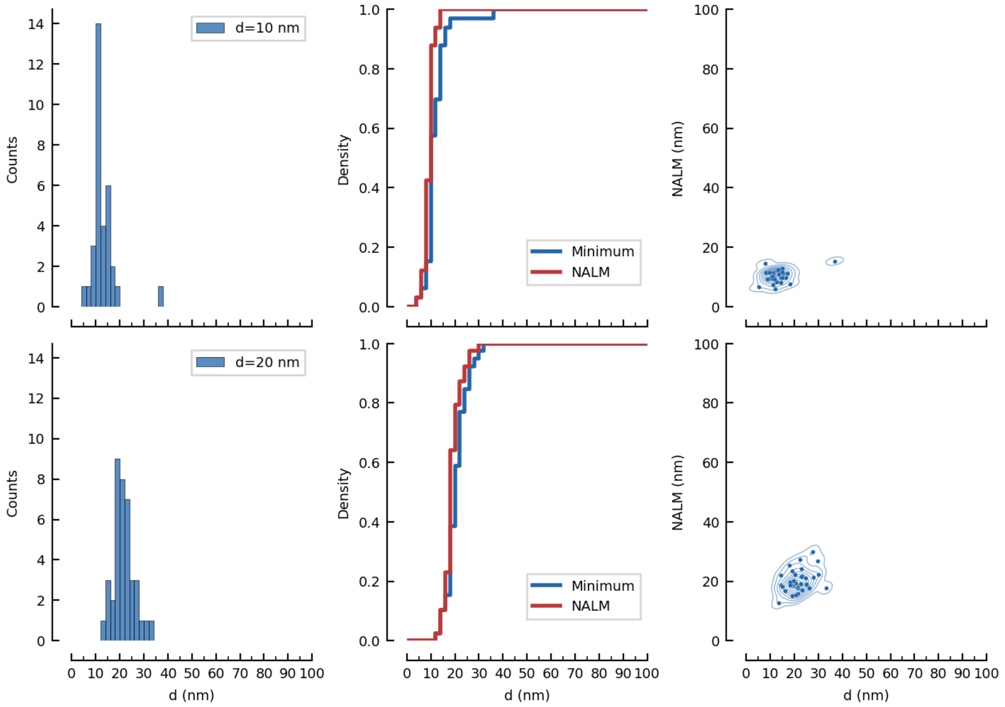

Figure 10: Statistical information on the distance estimation from line scans as described above for 10 nm and 20 nm.

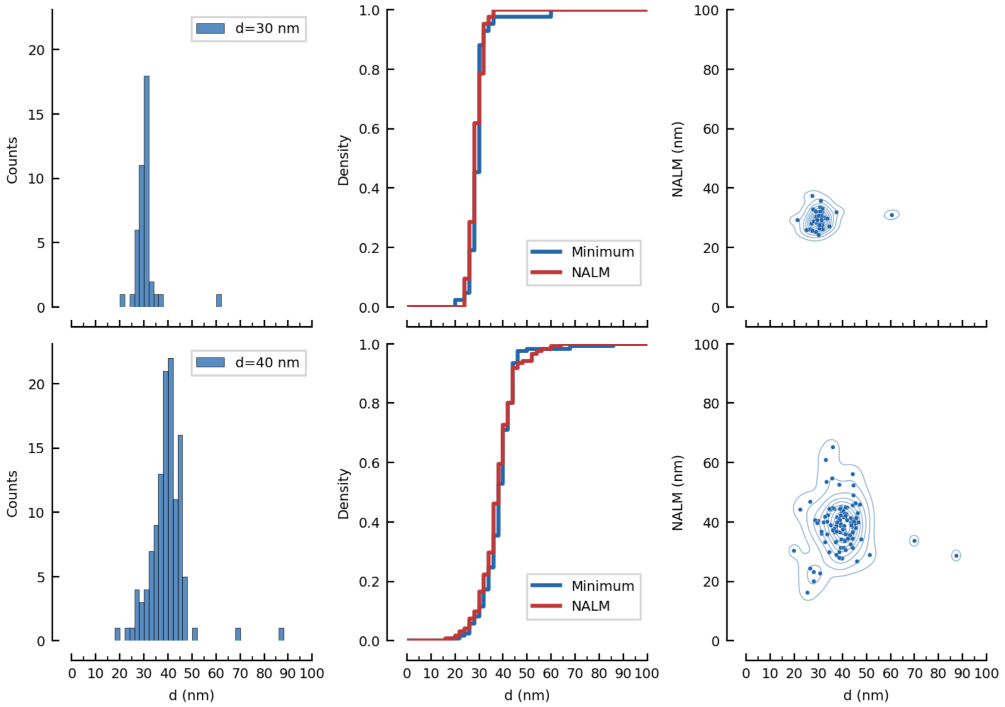

Figure 11: Statistical information on the distance estimation from line scans as described above for 30 nm and 40 nm.

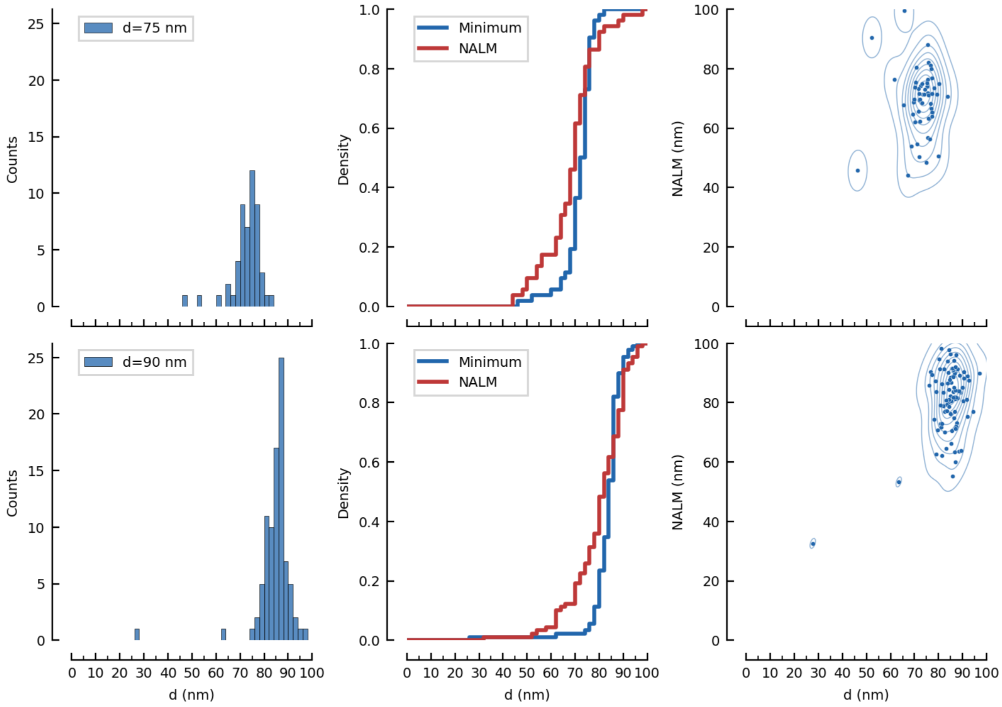

Figure 12: Statistical information on the distance estimation from line scans as described above for 75 nm and 90 nm.

#### 2.3 Control Method: Bleaching Steps

Recording line scans of a Nanoruler until both molecules are bleached offers an elegant way to run a control method for the distance estimate. Since it is very unlikely that both molecules bleach at the same time, a full trace contains a two molecule segment and a single molecule segment. While the minimum of the line scan is located at the centre-of-mass of the system when both molecules emit, it jumps to the remaining molecule once the other has bleached. Hence, the centre-of-mass shift is a convenient way to have an independent control for the distance estimate. However, it relies on the presence of a bleaching step and offers only an endpoint-like analysis, but no access to dynamic changes in distance with respect to time. This so called *Nanometer-localized multiple single-molecule fluorescence microscopy* has also been introduced in [4].

By employing a simple polynomial model that is fitted to the minimum, we extract the position of the minimum at any given time, average over the center-of-mass positions of the two molecule trace and do the same for the single molecule trace after the first bleaching step (see Figure 13. The molecule distance can be estimated as twice the lateral shift of the minimum.

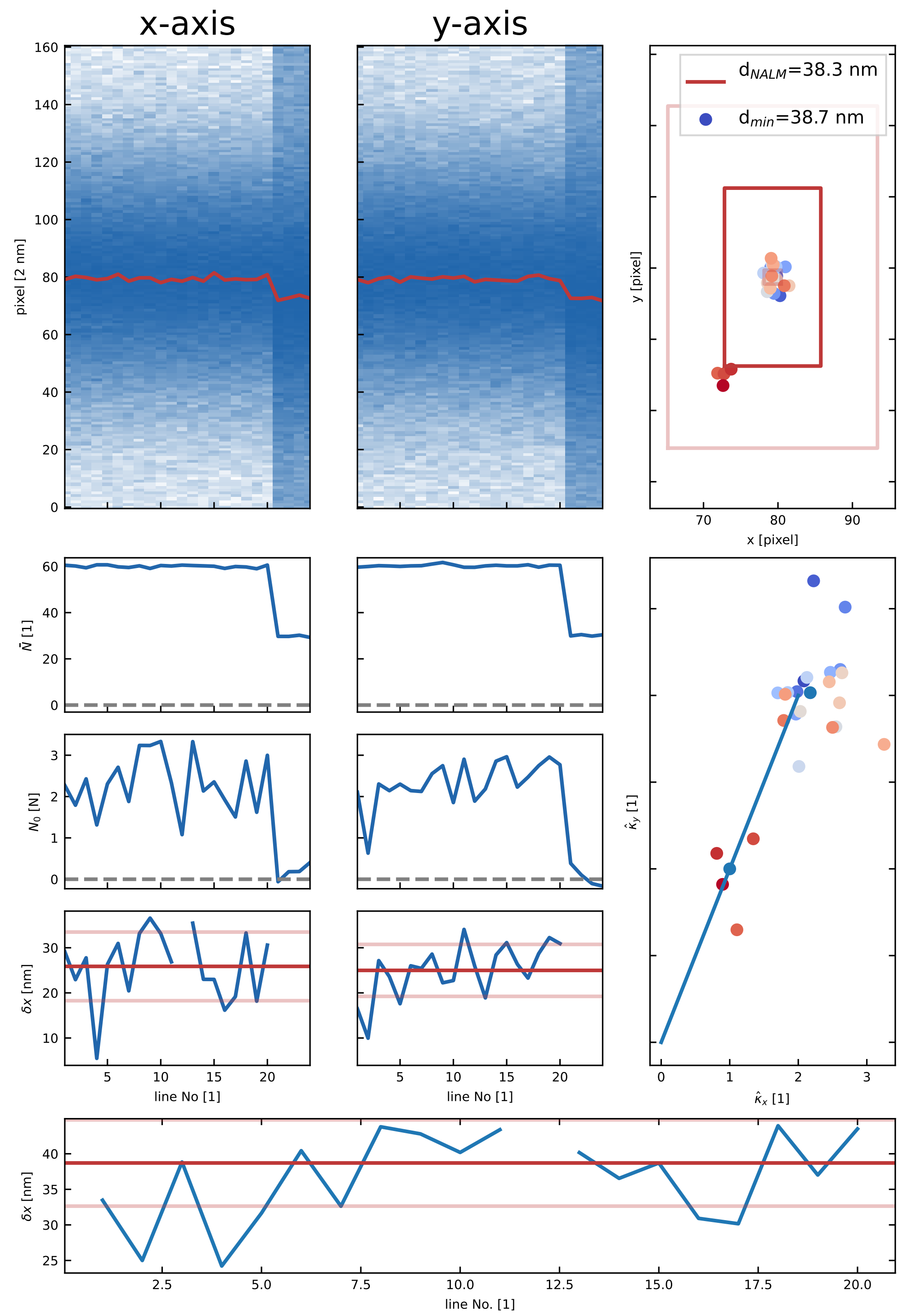

Figure 13: Extracting molecule distances via center-of-mass shifts due to bleaching steps for a 40 nm ruler. First row: counts with respect to scanning position and time (line). The center-of-mass position is marked with red and clearly shifts after the bleaching step. The positions obtained from the distance estimate with the minimum are plotted as scatters in the right plot, together with the distance estimates from the control for each axis. Panels below: counts in the minimum and estimated distances for each axis. Right spanning panel in middle row: estimated curvatures for each line. Lowest panel: norm of distance estimate wrt time (line number).

### 3 MINFLUX Analysis

#### 3.1 Data analysis

We leverage an iterative MINFLUX-like approach in order to sample the two emitters only in a region around the minimum of the illumination pattern (see [5]). In contrast to a full $2\pi$ line scan, this drastically reduces the number of photons collected, increases the speed of the measurement and ensures the collection of the most informative photons. As before, we only vary the phase of the illumination pattern and keep the spatial position of the envelope[3].

While the first iterations serve to localize the centre-of-mass of the system (position of the minimum), only photons of the last iteration are used to estimate the sources' separation. In the last iteration, we sample the system at three positions: $\varphi_- = x_0 - L/2$, $\varphi_0 = x_0$, $\varphi_+ = x_0 + L/2$, where $x_0$ denotes the centre-of-mass of the system and is updated after each measurement triple. This procedure is repeated consecutively in each axis, such that we record one x-triple, one y-triple and so forth. All measurement positions and counts form one trace. Since we are able to separate the two sources from their cumulative signal, we do not have to wait for bleaching steps. This would take too long and expose the measurement to ambient noise such as vibrations, temperature fluctuations or lateral drifts.

After the measurements, the trace is split into the x- and y-axis, which results in three point clouds at $\varphi_-$, $\varphi_0$, $\varphi_+$. Since the molecules are subject to a plethora of photophysical effects and especially a change of brightness would alter the distance estimate, we segment the trace with a hidden Markov-Model to account for these changes and treat the different segment separately.

Trace segmentation and correction for change in brightness.

The different segments are fitted with a polynomial estimator as described in subsection 1.4. This yields the parameters $a_0$ and $a_1$ of Equation 25, which allows to calculate the distance of the sources. We use a bootstrapping approach to bin the measurement tuples by different photon numbers or time-bins. This allows to extract subsequent distance estimates on a single Nanoruler.

Within each measurement campaign, a calibration measurement is performed to extract the initial visibility $\nu_0$. This can vary depending on the performance of the instrument. Since the distance estimate is very sensitive to the initial fringe contrast, we calibrate the analysis to the actual performance of the instrument in that session. A background estimate is performed on the same sample where we measure a batch of Nanorulers. Hence, we do a global background correction instead of a local (per Nanoruler) background estimate as is the case for the line scans.

[3] For more details on the phase-scanning MINFLUX approach see [1]

### 4 Multiple Emitters

We now touch on the parameter estimation for more than two point scatterers. Up to now, our formalism includes two scatterers, but in principle the concept should also apply to more than two scatterers. Here, we propose a protocol which allows to estimate positions of more than two sources simultaneously with an intensity minimum. We investigate the estimation capabilities of this concept by means of Monte-Carlo simulations. It turns out, that the advantage of the minimum remains for moderate spatial extent of an ensemble of emitters, and that an increasing emitter density also increases the resolution capability Figure 14.

#### 4.1 Monte-Carlo Simulation for geometrically constrained problems

In order to give a figure of merit for systems of more than two molecules, we first constrain their possible spatial arrangements to certain geometrical shapes, parametrized by a single parameter $d$. For a linear arrangement, $d$ denotes the nearest-neighbour distance of the sources. For a regular polygon $d$ is the diameter of the circle circumscribing all sources. For a grid-like structure $d$ denotes the grid constant. We perform a Monte-Carlo simulation of measurements for the aforementioned spatial arrangements with varying number of emitters and for a range of the scaling parameter $d$. We compute the relative error as the ratio of the Root-Mean-Square-Error (RMSE) of the estimated distance and the ground truth distance $RMSE(\langle d\rangle)/d$. Interestingly, even for more than two sources the parameter $d$ can be estimated precisely, which is not possible if a diffraction maximum is used. Hence, the advantage of employing intensity minima remains in the presence of more than two emitters. We note that increasing the emitter density allows for a more precise determination of the scaling parameter $d$ (Figure 14). This is not trivially anticipated, keeping in mind the problem of single parameter estimation when using diffraction maxima. However, further work is necessary to fully unfold the potential of this method and characterize its limitations.

#### 4.2 Cramer-Rao-Bound for multiple emitters in 1D

For multiple emitter arranged on a line, we can calculate an analytical expression for the Cramer-Rao-Bound. Assuming a negligible background and three probing positions at $\pm L/2, L = 0$, we get an uncertainty of

$$\begin{aligned}\sigma_{CRB} =&2\left(-\frac{\sin^2\left(\frac{d}{2}\right)\left(\sqrt{\frac{\cos(dm)-1}{\cos(d)-1}}-m\right)\left(2\cos\left(\frac{L}{2}\right)\left(\sqrt{\frac{\cos(dm)-1}{\cos(d)-1}}+m\right)+\sqrt{\frac{\cos(dm)-1}{\cos(d)-1}}-5m\right)}{N\left(-\cos\left(\frac{L}{2}\right)\sqrt{\frac{\cos(dm)-1}{\cos(d)-1}}-(\cos(L)+1)\sqrt{\frac{\cos(dm)-1}{\cos(d)-1}}+m(\cos(L)+1)+m\right)}\times\right.\\ &\left.\times\frac{\left(\cos\left(\frac{L}{2}\right)\sqrt{\frac{\cos(dm)-1}{\cos(d)-1}}-m\right)}{\left(m\cos\left(\frac{dm}{2}\right)-\cot\left(\frac{d}{2}\right)\sin\left(\frac{dm}{2}\right)\right)^2}\right)^{1/2}\end{aligned} \tag{93}$$

for the estimate of $d$.

In the limit $L \to 0$, we find the simpler expression

$$\sigma_{CRB} = \frac{2\left|\frac{\left(m-\sqrt{\frac{\cos(dm)-1}{\cos(d)-1}}\right)\sin\left(\frac{d}{2}\right)}{m\cos\left(\frac{dm}{2}\right)-\cot\left(\frac{d}{2}\right)\sin\left(\frac{dm}{2}\right)}\right|}{\sqrt{N}}. \tag{94}$$

However, we note that an infinite probing range L is not necessarily the best option to achieve the most precise estimate for more than two emitters. In fact, we observe that there is an ideal value for the probing range depending on the number of emitters.

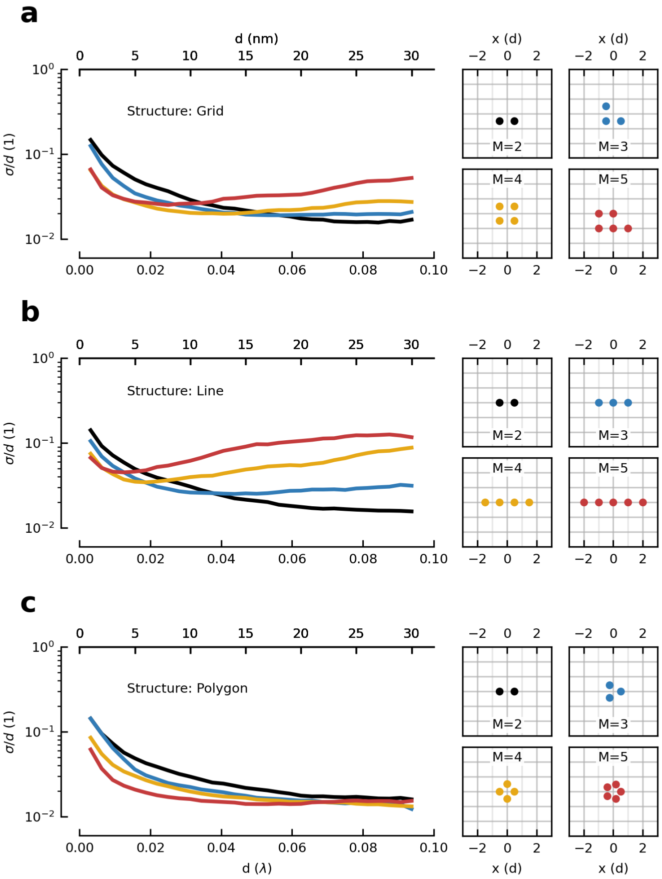

Figure 14: Breaking the diffraction-limit for more than two sources. Numerical simulation of the relative co-localization error for multi-emitter ensembles at N=500 collected photons. The advantageous scaling of the relative error remains valid for more than two sources. All systems are constrained to a single scaling parameter d. Its relative error is computed as $\sigma/d = \mathrm{RMSE}(d)/d$. a: Relative error for M emitters arranged on a grid with grid constant d in each axis (see cartoons) . At small distances, a higher number of sources can be resolved more precisely than only two. b: Relative error for M emitters arranged along a line at recurring distance d. Again, at small distances, a higher number of sources can be resolved more precisely than only two. For larger distances, the addition of another emitter comes at the cost of entering regions of higher intensity and losing the advantage of the intensity minimum. c: Relative error for M emitters placed on a circle of diameter d, forming a regular polygon. In this case, adding a source does not change the spatial extent of the ensemble, but increases the emitter density. This leads to an advantage of multiple sources compared to few over a larger range of the separation parameter d.

### 5 Sample Preparation

#### 5.1 Preparation of DNA-Origamis/Nanorulers

##### 5.1.1 Cleaning Process

To clean the cover slips (170 $\mu$m, No 1.5H, Paul Marienfeld GmbH & Co. KG, Lauda-Königshofen, Germany) measuring 15-18 mm in width, the following procedure was followed. Initially, they were gently wiped using a lint-free cloth sprayed with acetone. Subsequently, another cloth sprayed with Isopropanol was used to wipe them. After rinsing the cover slips with Isopropanol, they were dried using a nitrogen flow. Finally, a plasma cleaning process (using oxygen) was carried out for 5 minutes at 200 W.

##### 5.1.2 Construction of Flow Chamber

To facilitate the easy exchange of incubation solutions during preparation and buffer changes between experiments, flow chambers were created. This involved attaching two narrow strips of double-sided tape (Scotch Double Sided Tape, 3M, Saint-Paul, USA) to an object carrier, forming a channel approximately 5 mm wide. A cleaned coverslip was then placed on top of the tape and secured by applying gentle pressure.

##### 5.1.3 Preparation of DNA-Origamis/Nanorulers

The following reagents are required for the procedure:

1. BSA-Biotin
2. Streptavidin
3. Biotin functionalized - DNA-Atto 647N Nanoruler (Gattaquant)
4. Imaging buffer:
    - PBS
    - Glucose
    - Trolox
    - Methylviologen
    - Pyranose Oxidase
    - Catalase

The following steps were followed to prepare Nanorulers of various sizes:

1. Clean Coverslip (see subsubsection 5.1.1)
2. Prepare flow channel (see subsubsection 5.1.2)
3. Pipette the following through the flow channel:
    - 200 $\mu$L PBS (flushing the flow channel)
    - 15 $\mu$L BSA-Biotin (1:2 in PBS), wait two minutes
    - 200 $\mu$L PBS (flushing the flow channel)
    - 15 $\mu$L Strepatvidin (1:2 in PBS), wait two minutes
    - 200 $\mu$L PBS (flushing the flow channel)
    - 15 $\mu$L GattaQuant Rulers (diluted 1:50 or 1:100 in PBS), wait two minutes
    - 600 $\mu$L PBS (flushing the flow channel)
    - 15 $\mu$L Buffer
4. Seal edges with epoxy
5. Store in fridge

The buffer consists of an imaging buffer (IB), Methylviologen (MV) and an enzyme system. It is prepared as follows: mix 100 $\mu$L IB, 1 $\mu$L MV and 1 $\mu$L enzyme system. Reagents for the buffer:

- Imaging Buffer:
  - PBS (basis)
  - 10% Glucose (wrt mass)
  - 2 mM Trolox (wrt final concentration in IB)
- 0.2 M Methylviologen
- enzyme system: 10 mg Pyranase Oxidase + 170 $\mu$L PBS + 80 $\mu$L Catalase

To facilitate productive sample preparation, it is helpful to keep the following components as freezed stocks:

- BSA-Biotin (0.5 mg/ml in PBS) in 20 uL alliquots
- Streptavidin (0.5mg/ml in PBS) in 20 uL aliquots
- Buffer components:
  - IB
  - MV
  - enzyme system
