## Supplementary figures and images for "Diffraction minima resolve point scatterers at tiny fractions (1/80) of the wavelength"

### Supplementary Video

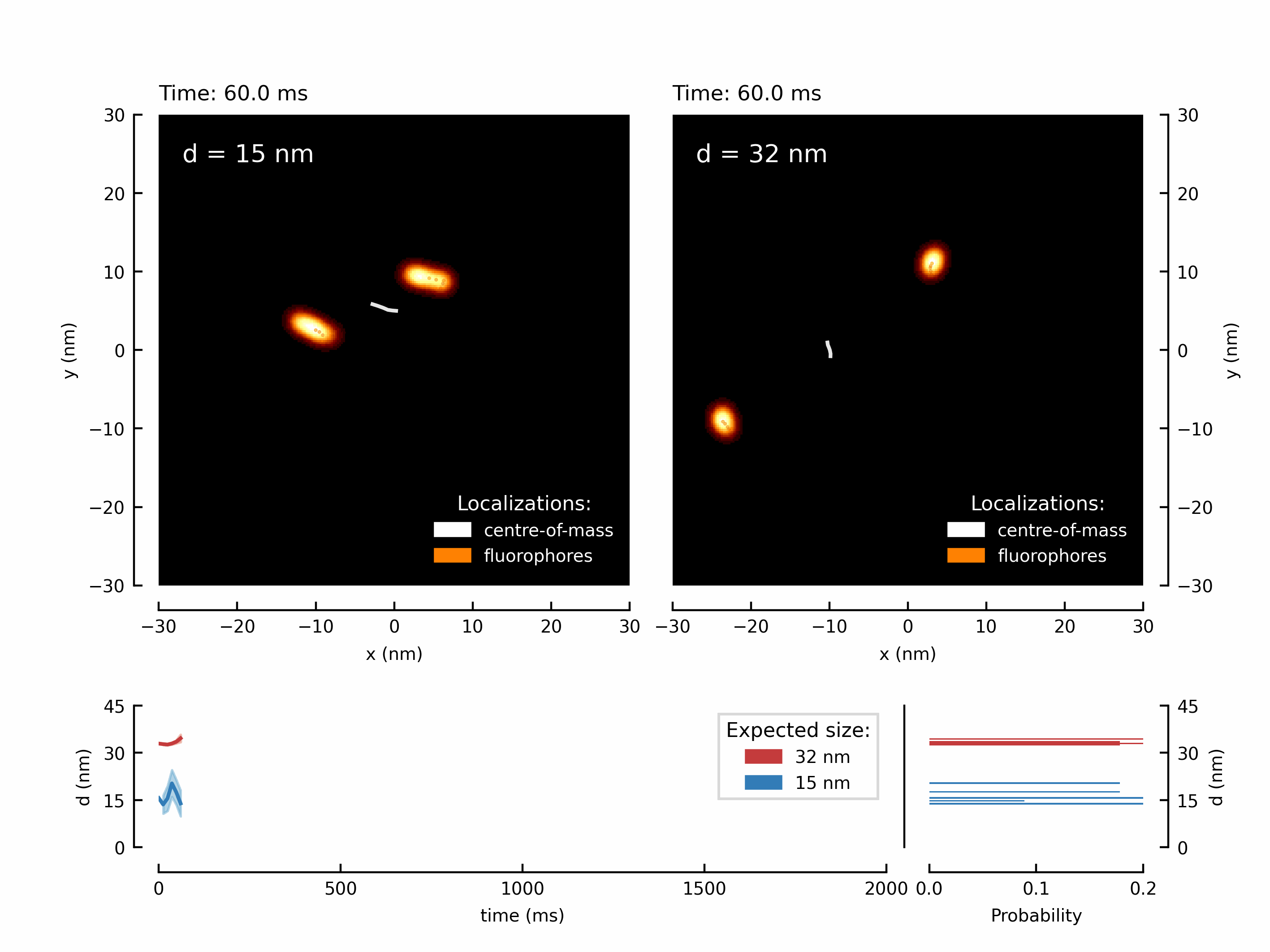
